# Membrane phosphoinositides allosterically tune β-arrestin dynamics to facilitate GPCR core engagement

**DOI:** 10.1101/2025.06.06.658200

**Authors:** John Janetzko, Jonathan Deutsch, Yuqi Shi, Dirk H. Siepe, Matthieu Masureel, Weijing Liu, Rosa Viner, Asuka Inoue, Steven Chu, Brian K. Kobilka, Rabindra V. Shivnaraine

## Abstract

Arrestin proteins bind active G protein-coupled receptors (GPCRs) through coordinated protein-protein, protein-phosphate, and protein-lipid interactions to attenuate G protein signaling and promote GPCR internalization and trafficking. While there are hundreds of diverse GPCRs, only two β-arrestin isoforms (βarrs) must recognize and engage this wide range of receptors with varied phosphorylation patterns. Traditional models suggest that βarr activation requires displacement of its autoinhibitory C-tail by a phosphorylated GPCR C-terminus; however, this paradigm fails to explain how minimally phosphorylated GPCRs still complex with βarrs. Using single-molecule Förster resonance energy transfer imaging and hydrogen-deuterium exchange mass spectrometry, we observe basal dynamics in which the βarr1 C-tail spontaneously releases from the N-domain, transiently adopting an active conformation that can facilitate binding of the phosphorylated GPCR C-terminus. We further demonstrate the importance of an intermediate state of βarr1 arising from spontaneous C-tail release stabilized by the membrane phosphoinositide PI(4,5)P_2_. Both PI(4,5)P_2_ and mutations in the proximal or middle regions of the C-tail shift βarr1 towards a partially released state, revealing an allosteric connection that informs a refined model for βarr activation. In this model, membrane engagement conformationally primes βarrs prior to receptor binding, thereby explaining how βarrs are recruited by diverse GPCRs, even those with limited C-terminal phosphorylation.

## INTRODUCTION

Two β-arrestin isoforms (βarrs) (βarr1 and βarr2) serve as multifunctional scaffolding proteins that shape the signaling outcome of hundreds of G protein-coupled receptors (GPCRs)^1,2^. Agonist binding promotes G protein engagement and leads to phosphorylation of intracellular loops and/or the C-terminus by various kinases, including GPCR kinases (GRKs)^3^. Through the action of these kinases, distinct phosphorylation patterns direct the interaction with βarrs^4,5^. Importantly, these different phosphorylation patterns can promote distinct βarr conformations and lead to divergent signaling outcomes^6–8^. Once bound to the receptor, βarrs desensitize G protein signaling^9^, promote internalization via clathrin-mediated endocytosis^9^, and modulate the interaction between GPCRs and other signaling effectors^9^.

For βarrs to engage the core of a GPCR, they must first transition from a basally autoinhibited state to an active conformation^10^. The primary gating mechanism for this process is thought to be the release of the βarr C-terminal tail (C-tail), which is displaced by the GRK-phosphorylated GPCR C-terminus (Rpp tail) which in turn binds to the N-lobe of βarr^11^. The C-tail of βarr1 and 2 also contains binding sites for components of the endocytic machinery (i.e., clathrin and adaptin proteins) that are exposed following C-tail release^12,13^. However, the sequence of events leading to a fully assembled GPCR-βarr complex has remained unclear. Further, understanding how assembly occurs is complicated by the structural diversity of GPCR-βarr complexes. With only two βarrs to regulate hundreds of GPCRs, βarrs must accommodate a wide range of phosphorylated epitopes and orientations. Several GPCR-βarr structures have been solved, revealing snapshots of the diversity within these complexes^14–24^. These structures reveal a range of binding modes for the engagement of a GPCR by βarrs. The prevailing model is that assembly proceeds via the initial formation of a so-called “tail-engaged” state, where the βarr is bound only by a phosphorylated region of the GPCR and does not directly engage the transmembrane core (Figure 1A). From this state, the GPCR-βarr complex is understood to evolve into a more stable core and tail-bound (fully assembled) state^25,26^. However, molecular dynamics (MD) simulations have shown that the transmembrane core is able to stabilize active conformations of βarrs^10^ and “tail-free core”-bound states, have been described^27–29^, where the assembly is phosphorylation-independent, or proceeds with minimal GPCR phosphorylation. Yet, the present model fails to explain how activation of βarrs can proceed in these cases.

**Figure 1.**
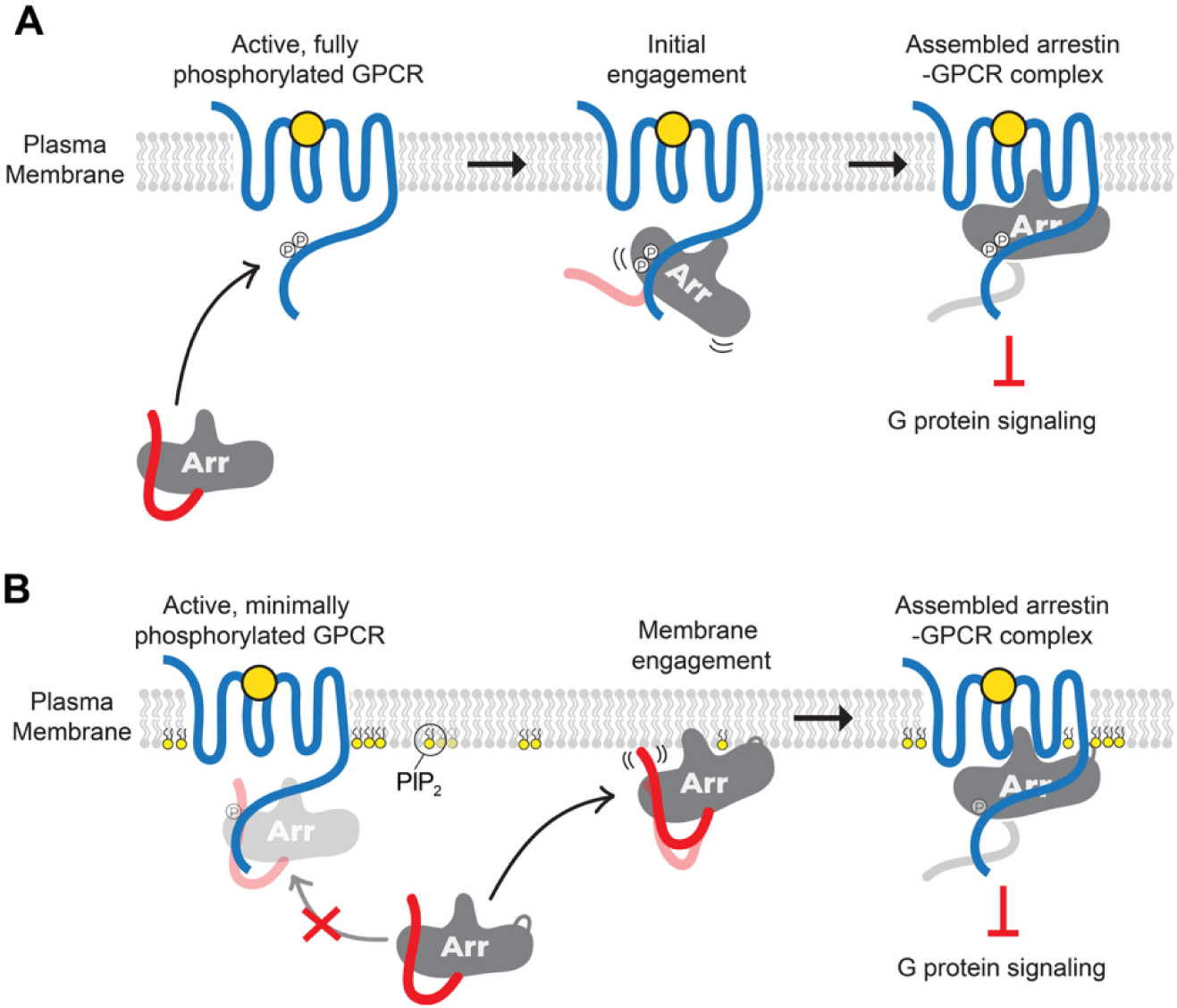
Models of β-arrestin assembly to GPCRs. Schematic of arrestin recruitment to an agonist-bound GPCR in the plasma membrane (PM). (A) For strong βarr interacting GPCRs (class B), the Rpp motif can effectively capture the transient membrane-associated βarr1 population without requiring pre-activation of βarrs by membrane PIPs. (B) In contrast, for weak βarr interacting GPCRs (class A), pre-activation by membrane PIPs may function to increase the likelihood of productive engagements with an active but minimally phosphorylated GPCR.

Recently, increasing attention has been given to the role of βarr-interactions with lipid membranes. We and others have shown that membrane phosphoinositides (PIPs) play an important role in GPCR-βarr complexes^25,30–35^. In particular, it has been seen that plasma membrane PIPs, such as phosphatidylinositol 4,5-bisphosphate (PIP_2_), can bind to βarrs with high-affinity and can stabilize them at clathrin-coated structures even in the absence of an associated GPCR^36^. PIPs bind in the concave face of the βarr C-lobe, as shown in the first structure of a non-visual arrestin-receptor complex^14^, where it appears to stabilize the βarr1-GPCR core-engaged state. Our previous work^25^ demonstrated that βarr recruitment to various GPCRs could be stratified into two distinct behaviors based on whether the βarrs required PIP_2_ binding, broadly aligned with the previously described class A/B typology^37^. For class B GPCRs that strongly interact with βarrs, phosphorylation motifs within the GPCR C-terminus are sufficient to recruit βarrs. In contrast, class A GPCRs, which have weaker phosphorylation-dependent interactions with βarrs, require the PIP_2_ binding site in βarr for effective engagement, desensitization, and internalization^25^.

Single particle tracking experiments in live cells recently revealed that βarrs may first transiently engage the plasma membrane prior to engaging a target GPCR^32^. Importantly, this study showed that while the PIP binding site of βarrs was not necessary for sampling the plasma membrane, it was necessary for effective GPCR engagement. βarrs have been shown to bind lipid membranes through additional motifs, including the C-edge (344 and 197 loops)^38^, and these loops were critical for transient sampling of the membrane by βarrs. This raises the question of when and how membrane PIPs mediate GPCR engagement, especially for GPCRs that lack strong phosphorylation motifs^39^.

Here, we used single-molecule Förster resonance energy transfer (smFRET) to monitor the dynamics of the βarr1 C-tail in its basal state as well as in complex with a phosphorylated GPCR, phosphopeptide mimics of a GPCR C-terminus, and membrane phosphoinositides. We also investigate how mutations in the βarr1 C-tail that destabilize the autoinhibited state of βarr1 alter the dynamics in the context of these inputs. This approach allowed us to directly observe transient dynamics and intermediate activation states that are largely inaccessible through ensemble or structural methods. We further integrate these findings with hydrogen-deuterium exchange mass spectrometry (HDX-MS) and other biochemical cell-based assays to obtain a refined picture of how βarr1 activation and differences in GPCR phosphorylation may alter the path of GPCR-βarr1 assembly. Our revised model demonstrates how C-tail conformational dynamics are allosterically tuned by membrane inputs such as PIP_2_ to mediate recruitment, especially to minimally- or non-phosphorylated GPCRs^40,41^ (Figure 1B).

## RESULTS

To probe the conformational changes of the βarr1 C-tail, we used a cysteine-free construct^42^ and inserted cysteine residues at positions 12 (on the N-terminal β-strand) and 387 (in the proximal C-tail)^25^ (see Methods) (Figure 2A). This construct, βarr1-12-387, was stochastically labeled with maleimide conjugates of Alexa 488 (donor) and ATTO647N (acceptor) fluorophores, which did not alter the monodispersity of the protein as shown with size-exclusion chromatography (SEC) (Figure S1A). WT-like functionality of this labeled βarr1 C-tail sensor, βarr1-12-387-AF488-AT647, has previously been demonstrated by its bulk FRET response to a phosphopeptide corresponding to the fully phosphorylated state of the human V2 vasopressin receptor C-terminus (V2Rpp)^43–45^, bearing 8 phosphorylated residues, as well as to a soluble derivative of the plasma membrane phosphoinositide PI(4,5)P_2_ (PIP_2_)^25^. For smFRET measurements, βarr1-12-387-AF488-AT647 was site-specifically biotinylated through an N-terminal Avi tag^46^ and specifically immobilized (Figures S1B and S1C) on passivated quartz slides via a Neutravidin-biotin bridge. Individual molecules of the βarr1 C-tail sensor captured on the surface were imaged using a prism-TIRF microscope, enabling real-time monitoring of the C-tail movements through anti-correlated changes in the donor and acceptor fluorescence and corresponding FRET efficiency (Figures 2A and 2B).

**Figure 2.**
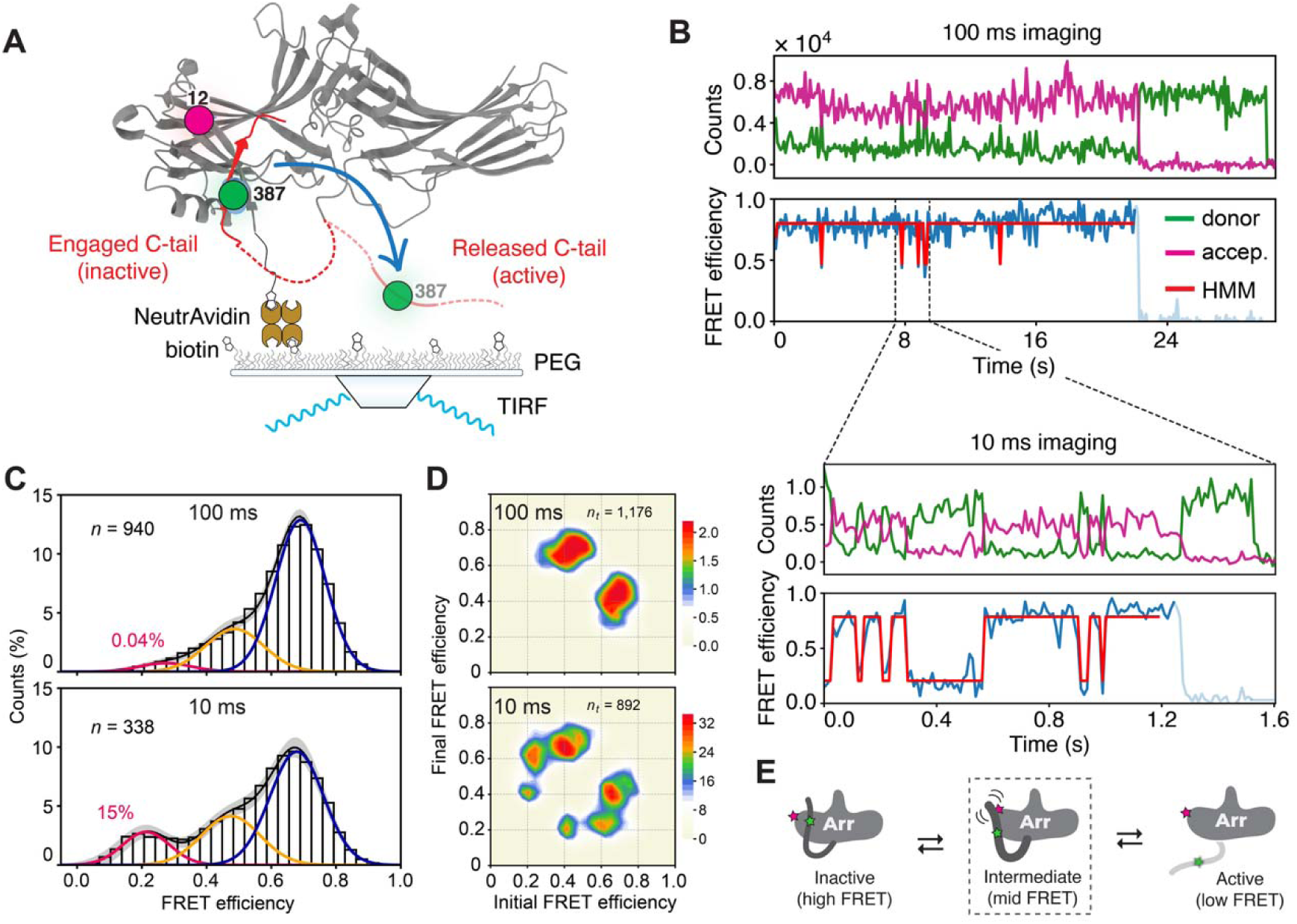
The βarr1 C-tail is dynamic in its basal state. (A) Schematic of the smFRET experiments with surface-tethered βarr1 labeled with fluorophores (donor in green; acceptor in magenta) shown on the inactive structure (PDB: 1G4M). (B) Example fluorescence and smFRET traces with state idealization (red line) for basal βarr1 recorded 100 ms (top) and 10 ms (bottom) imaging speeds. (C) Population FRET efficiency histograms and three-state GMM for basal βarr1 at 100 ms (top) and 10 (bottom) ms with different low-FRET (active) state occupancies (pink text). Error bands, the 95% confidence interval (c.i.) in the GMM fits calculated from 100 bootstrap samples of the *n* total FRET traces. (D) Transition density plots displaying the mean FRET values before (x-axis) and after (y-axis) each transition at 100 ms (top) and 10 ms (bottom). Scale bar, 10^-3^ transitions per bin per second; *n_t_* total transitions per condition. (E) Cartoon showing how smFRET efficiency is related to C-tail release. See also Figure S1 and S2.

### Basal dynamics of the βarr1 C-tail

Under basal conditions, where the C-tail is expected to reside in a docked position against the N-lobe^47^, smFRET measurements of surface-tethered βarr1 predominantly showed a high-FRET state (Figures 2B and 2C). However, we also observed rapid, transient excursions to lower-FRET states that we ascribe to spontaneous disengagement of the C-tail from the N-domain groove (Figures 2B, S1D and S1E). Population histograms of the smFRET efficiency recorded at both 100 ms and 10 ms integration times fit to a gaussian mixture model (GMM) identified three main FRET states centered at ∼0.21, ∼0.44 and ∼0.68 (Figure 2C). Regardless of imaging speed, the majority (>60%) of basal βarr1 occupied the inactive, high-FRET (∼0.68) state, representing the fully bound (i.e., auto-inhibited) position of the C-tail. Imaging at 10 ms, however, resulted in nearly a 4-fold increase in the fractional occupancy (*F*) of the low-FRET (∼0.21) state (Figure 2C), indicating better temporal resolution of the rapid C-tail fluctuations.

Consistent with this observation, idealization of the traces using a hidden Markov model^48^ (HMM) (see Methods) revealed approximately an order of magnitude more reversible, high-to-low FRET transitions at 10 ms compared to 100 ms, with rates of >3 s^-1^ and 0.25 s^-1^, respectively (Figure 2B and 2D). At 100 ms integration time, transitions primarily occurred between the high-FRET (∼0.68) state and a broad mid-FRET (∼0.4-0.5) state, whereas at 10 ms, transitions were observed between all three FRET states of the C-tail (Figures 2D and 2E). The mean dwell time (τ) of the better-resolved, low-FRET excursions was ∼100 ms, making these events difficult to cleanly resolve at the 100 ms per frame imaging speed^49^. Therefore, rapid (<300 ms) transitions of the C-tail from the high- to the low-FRET states may manifest as a broadened mid-FRET state due to time-averaging between the extrema states (Figure 2B).

Analogous measurements using ATTO643 (a hydrophilic variant of ATTO647N) as the FRET acceptor yielded similar ensemble V2Rpp binding (Figure S2A) as well as comparable basal C-tail dynamics at 10 ms (Figure S2B-D). These results suggest that the observed transitions reflect true C-tail conformational changes rather than photophysical artifacts or environment interactions of the probe. Such spontaneous and short-lived transitions to a released C-tail conformation may explain previous observations that apo βarr1 can transiently bind the active-state-specific binder Fab30^25^. Together, these data indicate that the proximal C-tail of βarr1 is intrinsically dynamic, even in its basal state.

### Modulation of βarr1 C-tail dynamics by GPCR phospho-tail mimetics

The low- and mid-FRET basal states observed for βarr1 represent disengaged conformations of the C-tail, which may facilitate the binding of the Rpp tail. To examine the effect of a phosphorylated GPCR C-terminus on C-tail dynamics, we imaged the βarr1-12-387-AF488-AT647N C-tail sensor in the presence of V2Rpp (Figure 3A). In measurements of surface-tethered βarr1 taken at 10 ms, V2Rpp binding resulted in a concentration-dependent increase in the low-FRET (∼0.21) state and a corresponding depletion of the high-FRET (∼0.68) inactive state (Figures 3A, S2E, S2F and S3A). The EC_50_ of this V2Rpp-induced low-FRET occupancy was 11.8 ± 1.6 μM (Figure 3B), in line with previous measurements of V2Rpp binding to βarr1^4,5,45^. To confirm that these measurements were consistent with the effects of a full-length receptor, agonist-bound (NTS8-13) and GRK5-phosphorylated human neurotensin receptor type 1 (NTSR1)^14^ was introduced into an imaging chamber containing immobilized βarr1. Saturating concentrations of phosphorylated NTSR1 (5 μM) and V2Rpp (62.5 μM) yielded nearly identical FRET distributions (Figures S3A and S3B).

**Figure 3.**
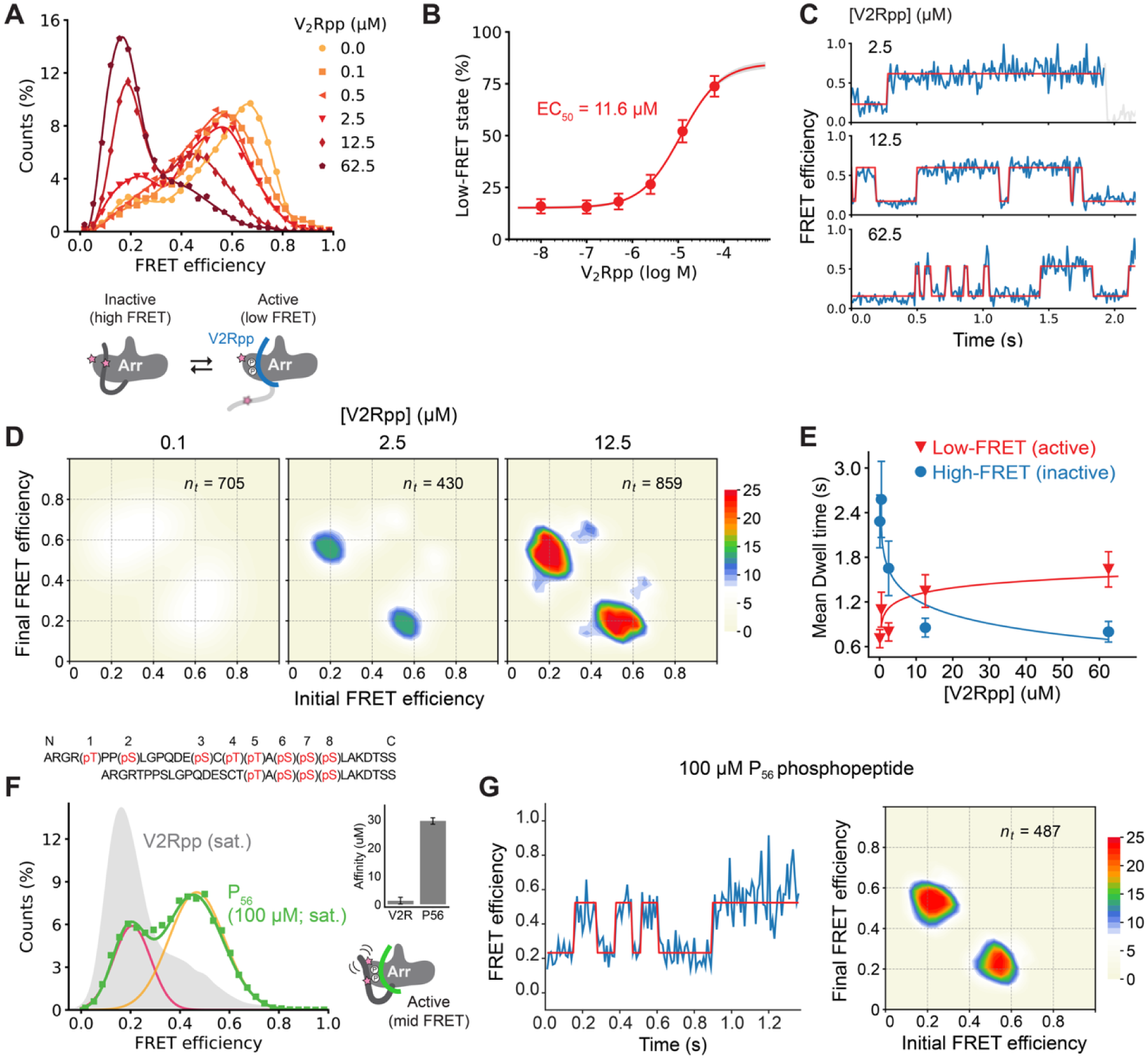
Receptor phospho-tail mimics differently activate the βarr1 C-tail. (A) Top: population FRET efficiency histograms (symbols) and b-spline fits (lines) for surface-tethered βarr1 in the presence of the indicated concentrations of V2Rpp. 1,715 molecules across all conditions. Bottom: cartoon showing a model of FRET changes induced by V2Rpp. (B) Ensemble average low-FRET (active) state occupancy (circles) induced by increasing concentrations of V2Rpp fit to a dose-response curve (Hill slope = 1; solid line) with an apparent EC_50_ value of 11.6 μM. Error bars, 95% c.i. of 100 bootstrap samples of the FRET traces. (C) Example smFRET traces (blue) with state idealization (red line) for the indicated V2Rpp concentrations. (D) Select transition density plots displaying the mean FRET before and after each transition (scaled to saturating V2Rpp). (E) Intensity averaged dwell time in the low-FRET (active) and high-FRET (inactive) states using a two-state HMM. Error bars, s.d. from 100 bootstrap samples of the FRET traces for each concentration. (F) Population FRET histogram of βarr1 imaged in saturating P_56_ shows an intermediate active state. Inserts: phosphorylation patterns (top) and binding affinities (right) for V2Rpp and P_56_ phosphopeptides. Error bars, s.e. of the regression estimate of the EC_50_. (G) smFRET trace (left) and transition density plot (right) for βarr1 in the presence of saturating (100 μM) P_56_. Scale bar, 10^-3^ transitions per bin per second (*n_t_* total transitions); same scale as in (D). All data is at 10 ms integration time. See also Figures S3 and S5, as well as Data S1.1.

Individual trajectories of the βarr1 C-tail revealed frequent (≤2.5 s^-1^) reversible transitions from the high- to the low-FRET state in the presence of V2Rpp (Figure 3C). Modeling the kinetics of these transitions showed that increasing concentrations of peptide reduce the dwell time of the high-FRET state by up to three-fold, resulting in a concentration-dependent increase in dynamics (Figures 3C-E, S3A, and S3C). These results are consistent with competition between V2Rpp and the βarr1 C-tail. In contrast, the low-FRET dwell time was prolonged in the presence of V2Rpp (Figure 3E), reflecting a mixture of two decay processes: slower (<1 s^-1^), first-order V2Rpp unbinding and the faster (>10 s^-1^) intrinsic C-tail dynamics present in the unbound βarr1 faction. The addition of Fab30, an antibody fragment raised against the V2Rpp-bound conformation of βarr1^43^, further stabilized the low-FRET state, increasing the dwell time by an order of magnitude (5 s versus 0.45 s) (Figures S3D and S3E). These data suggest that Fab30 dramatically reduces V2Rpp exchange by slowing its dissociation from βarr1, as observed previously^25,43,45^.

In addition to V2Rpp, we also assessed the effect of phosphopeptide composed of a sub-maximal phosphorylation pattern on the C-tail conformation and dynamics. Previous MD simulations showed that a V2Rpp-derived phosphopeptide, P_56_, lacking four N-terminal phosphorylation sites (Figure 3F, upper insert), was only able to promote a partial C-tail displacement^4^. In the fully phosphorylated V2Rpp, these phosphosites contact residues in the N-lobe and at the base of the βarr1 finger loop^43^. In bulk, the binding affinity of P_56_ to βarr1 was ∼30 μM, an order of magnitude weaker than the fully phosphorylated V2Rpp (EC_50_: ∼1.5 μM), based on FRET changes (Figure S3F). Moreover, displacement of the βarr1 C-tail with saturating P_56_ resulted in a higher tail asymptote in bulk FRET experiments compared to V2Rpp (∼0.3 versus ∼0.2; Figure S3F), indicating less net displacement of the βarr1 C-tail. Consistent with these bulk FRET results, smFRET measurements of βarr1-12-387-AF488-AT647 at a saturating concentration (100 μM) of P_56_ predominantly shifted the C-tail to the mid-FRET (∼0.44) state, rather than the low-FRET (∼0.21) state (Figures 3F and S3G). Additionally, we observed only partial (∼30%) occupancy of the low-FRET, approximately double the basal level (i.e., without phosphopeptide) (*F*_apo_ ≈ 0.16). Inspection of individual trajectories revealed reversible mid-to-low transitions (Figures 3G), occurring with similar (∼1 s^-1^) frequency to those observed with saturating V2Rpp (cf. Figure 3D). However, the average duration of the low-FRET dwells was shorter with P_56_ compared to V2Rpp, 0.5 s and 1.6 s, respectively. This suggests P_56_-bound βarr1 primarily adopts a partially activated (mid-FRET) conformation of the C-tail, likely because the incomplete phosphorylation pattern is not sufficient to fully release the C-tail and as such does not significantly stabilize transitions to the low-FRET state beyond the basal level (τ_apo_: ∼0.3 s). Based on this, we anticipate other phosphorylation patterns differ in their ability to release the βarr1 C-tail, stabilizing distinct proportions of the active states or giving rise to additional C-tail conformations not observed here.

### Effect of phosphoinositides on C-tail release

We and others have found that membrane phosphoinositides (PIPs) can regulate βarr activation and GPCR recruitment through their binding to the concave face of the C-lobe, away from the C-tail^25,30,35^. Recent ensemble FRET^25^ and NMR spectroscopy^33^ studies have suggested that a soluble derivative of phosphatidylinositol 4,5-bisphosphate (henceforth PIP_2_) can partially activate βarr1 in the absence of a GPCR, resulting in conformational changes in the C-tail without fully releasing it. Importantly, this effect is lost upon mutation of the three primary residues in the βarr1 C-lobe that contact the phosphoinositide head group (K232Q/R236Q/K250Q, henceforth 3Q)^25,30^. This effect of mutation on binding was confirmed by fluorescence polarization (FP) measurements using a BODIPY-derivative of PIP_2_, which showed that βarr1 3Q essentially lost binding to PIP_2_ (Figure 4A). Together, these findings demonstrate allosteric coupling between the βarr1 C-lobe and the C-tail.

**Figure 4.**
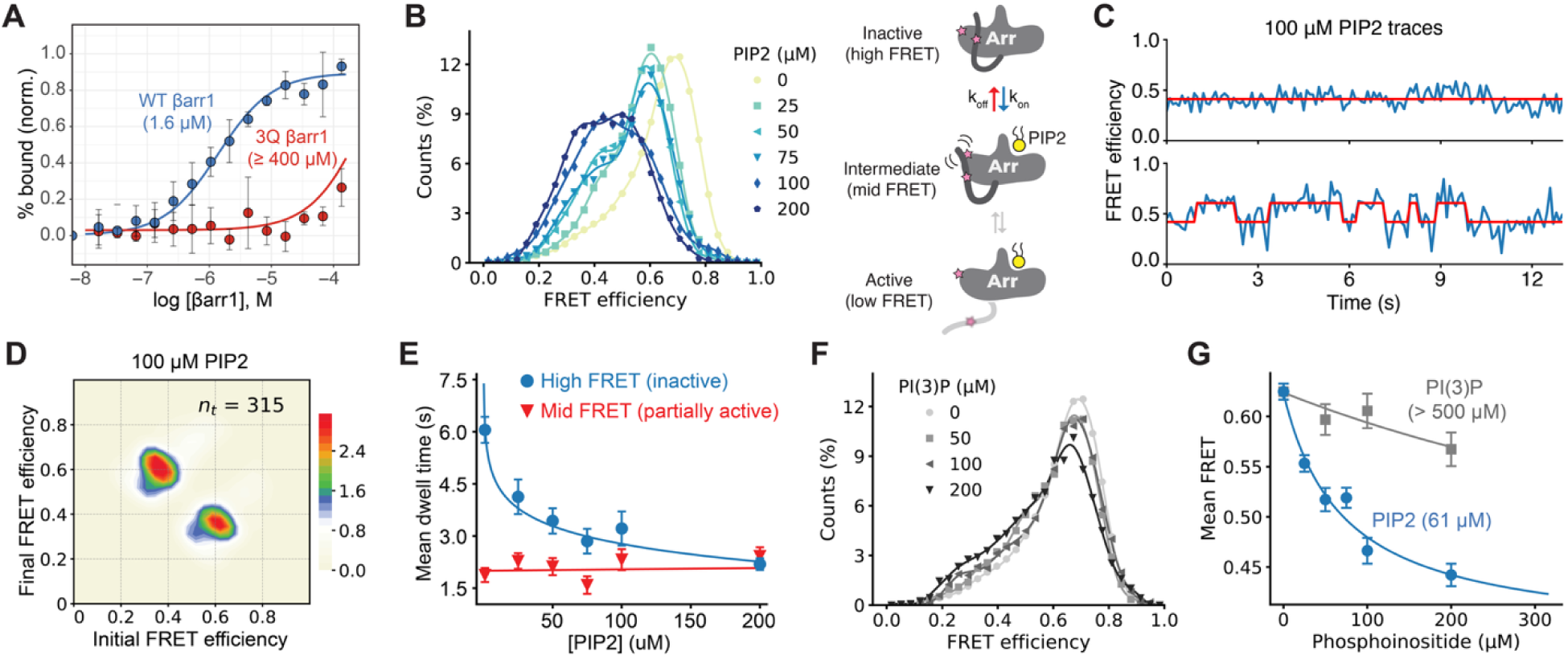
PIP_2_ binding releases the C-tail to an intermediate state. (A) Binding of BODIPY-PIP_2_ to WT or 3Q βarr1. Data are mean fluorescence polarization ± 95% c.i. (n = 5). Lines are fits to dose-response functions with EC_50_ of 1.6 μM for WT and > 400 μM for 3Q (assuming the same maximal binding). (B) Left: Population FRET efficiency histograms (symbols) and b-spline fits (lines) from imaging surface-tethered βarr1-12-387-AF488-AT647N at 100 ms imaging speed in the presence of the indicated concentrations of PIP_2_. N = 3,273 total molecules across all conditions. Right: Cartoon showing model of FRET changes induced by PIP_2_. (C and D) (C) Example smFRET traces and (D) transition density plot from experiments with 100 μM PIP_2_. (E) Mean dwell times (symbols) of the βarr1 C-tail in the high-FRET, inactive state (blue) and the mid-FRET, partially-displaced state (red) fit to a log-linear model (lines). Error bars, s.d. of 100 bootstrap samples of the FRET traces. (F) Population FRET histograms (symbols) and spline fits (lines) from experiments imaging βarr1 at 100 ms in the indicated concentrations of PI(3)P. N = 1,764 total traces across all conditions. (G) Ensemble average FRET efficiency induced by PIP_2_ (blue) and PI(3)P (gray). Lines are fits to dose-response functions with EC_50_ of 61 μM and > 500 μM for PIP_2_ and PI(3)P, respectively. See also Figure S4.

Measurement of the allosteric effects of PIP_2_ on the C-tail dynamics of surface-tethered βarr1-12-387-AF488-AT647 revealed a concentration-dependent redistribution from the high-FRET (∼0.68) state to a broadly distributed mid-FRET (∼0.44) state (Figures 4B-E and S4A-D). Both 100 ms measurements (Figures 4B and S4A) and 10 ms measurements (Figure S4B) showed similar results. To control for the impact of high (>100 μM) lipid concentrations on the fluorescent probes, we tested the effect of PI(3)P at similar concentrations, which showed an extremely weak affinity for βarr1^25,30^ (concentration-dependent shift in FRET efficiency, EC_50_ >700 μM; Figure 4F, 4G and S4E). In contrast, PIP_2_-induced changes in average smFRET had an EC_50_ of 58 ± 1.4 μM (Figure 4G), in line with prior ensemble measurements^25^. The effect on the C-tail conformation cannot be attributed to artifacts such as aggregation, as size-exclusion chromatograms of βarr1 in the presence of varying concentrations of PIP_2_ showed only a monomeric peak (Figure S4C).

Using hidden Markov modeling, we quantified the allosteric effects of PIP_2_ binding and dissociating on the transition rates of the C-tail between the high- and the mid-FRET states (Figures 4B-D). We observed prolonged (up to 15 s) dwells in the mid-FRET state, while HMM analysis revealed that the primary effect of PIP_2_ was reduction of the residence time of the high-FRET (inactive) state due to promoting transitions to the mid-FRET (∼0.18 s^-1^ at 100 μM; Figures 4C-E). The dwell times of the mid-FRET (“pre-activated”) state were not systematically affected (Figure 4E), indicative of concentration-dependent PIP_2_ binding and concentration-independent unbinding. Apparent rate constants derived from transitions observed at sub-EC_50_ concentrations of PIP_2_, were *k*_high-to-mid_ = 0.0025 uM^-1^ s^-1^ and *k*_mid-to-high_ = 0.48 s^-1^, for binding and unbinding, respectively (Figure S4D). However, direct measurements of PIP_2_ binding to βarr1 using radioligand^30^, or fluorescent BODIPY-derivatized PIP_2_ (Figure 4A), estimated an affinity of ∼2 μM, substantially less than the EC_50_ of the conformational changes (∼60 μM). This suggests that not every PIP_2_ binding event immediately shifts the C-tail to the intermediate state.

As the phosphopeptide and phosphoinositide binding sites are spatially distinct, this opens the possibility for synergistic inputs to βarr1. In experiments with saturating P_56_, addition of saturating PIP_2_ yielded an increase in low-FRET occupancy from 0.34 to 0.50 (Figures S5A and S5B). Crosstalk between P_56_ and PIP_2_ has a positively cooperative effect on the C-tail, where their co-exposure led to a low-FRET population (0.5 occupancy) that was approximately equal to the sum of each ligand alone (*F*_P56_ ≈ 0.34, Figure S3G; *F*_PIP2_ ≈ 0.17, Figure S4B). Individual trajectories showed transitions between the mid- and the low-FRET states with only rare (∼5 min^-1^) transient excursions to autoinhibited conformations with FRET_ideal_ > 0.6 (Figures S5C and S5D). Additionally, the C-tail was almost twice as dynamic in the presence of both ligands as it was with phosphopeptide alone (∼1.8 s^-1^ versus 3.1 s^-1^) (Figure S5D), likely due to the increased displacement that yields increased binding and unbinding events. These data suggest that when either P_56_ or PIP_2_ is bound to βarr1, it primarily adopts the partially engaged (mid-FRET) conformation with infrequent low-FRET transitions. When both ligands are bound, their interplay promotes transitions from the mid-to the low-FRET states, increasing the population of fully activated βarr1 (illustrated in Figure S5B). However, PIP_2_ was previously shown to have complex effects on the conformation of V2Rpp-bound βarr1, and for some weakly activating Rpp tails, crosstalk with PIP_2_ may be counteractive rather than cooperative^33^.

### Activating mutations and PIP_2_ synergistically release the C-tail

To further explore the molecular determinants of βarr1 activation and the role of the intermediate FRET state, we examined the effect of mutating either the proximal or middle regions of βarr1 C-tail. For the proximal C-tail mutation, we used the previously described F388A/V387C/I386A triple mutant (henceforth 3A) that anchors the proximal C-tail via hydrophobic interactions with leucine residues on α-helix 1 of the N-domain in what is referred to as the three-element interaction (3-EI) (Figure 5A)^50^. Specifically, 3A βarr1 has been shown to have decreased specificity between active and phosphorylated vs active but non-phosphorylated GPCR forms^50^.

**Figure 5.**
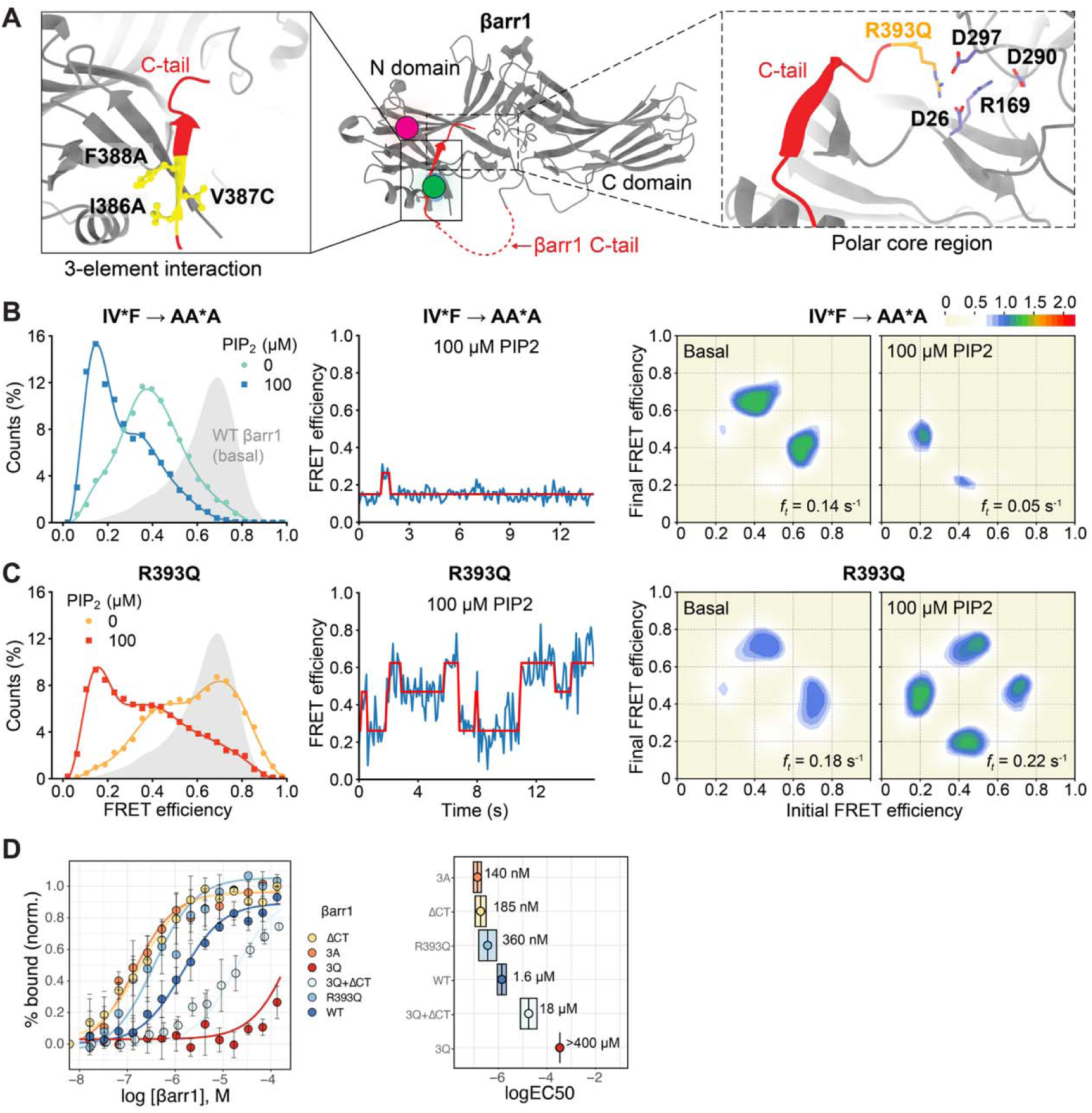
βarr1 C-tail mutants show enhanced response to PIP_2_. (A) Three-element (left insert) and polar core (right insert) interactions stabilize the autoinhibited state of the βarr1 C-tail (PDB: 1G4M). (B and C) smFRET experiments imaging (B) 3A*- and (C) R393Q βarr1 constructs in the presence and absence of 100 μM PIP_2_ at a 100 ms camera integration time. Left: population FRET histograms (symbols) with b-spline fits (lines); 692 and 635 total traces for 3A and R393Q, respectively. Gray density is WT βarr1 in the basal state. Middle: example smFRET traces in the presence of 100 μM PIP2. Right: Transition density plots in the absence (left) and presence (right) of 100 μM PIP_2_. Scale bar, 10^-3^ transitions per bin per second; *f_t_* denotes the overall transition frequency. The same scale is used as in Figure 2D, 100 ms. (D) FP measurement of saturation binding of BODIPY-PIP_2_ to various βarr1 constructs. Left: points are mean ± 95% c.i. (n = 5) of the normalized fluorescence polarization fit to dose-response functions (lines). Right: bars show EC_50_ values with 95% c.i. Indicated. For βarr1 3Q the bars are larger than the clipped axes due to the high EC_50_ and low confidence in the fit. Box shows best-fit EC_50_ values derived from regression estimates on left. See also Figure S5.

Single-molecule measurements of 3A βarr1 showed a nearly complete depletion of the high-FRET inactive state and instead indicate the C-tail primarily resides in an intermediate ∼0.4-0.45 FRET state at both 100 ms (Figures 5B and S6A) and 10 ms (Figure S6B) camera integration times. Because of the similarity between data recorded across both time regimes, further measurements of C-tail mutants were taken at 100 ms. In contrast to the parent βarr1 construct, treatment of the basally mid-FRET 3A mutant with 100 μM PIP_2_ caused a shift from the mid- to the low-FRET state (Figures 5B). In comparison, the increase in fractional occupancy of the respective states promoted by PIP_2_ was similar for both the parent (Figure S4A) and 3A (Figure S6A): 0.2 to 0.5 and 0.3 to 0.6, respectively. Furthermore, 3A βarr1 exhibited similar but reduced dynamics compared to the parent sensor, with PIP_2_ further suppressing the frequency of transition between the low- and the mid-FRET states (Figures 5B, right panel). Together, these observations suggest that disrupting the 3-EI dissociates the proximal segment of the C-tail, likely while leaving the C-tail anchored through the distal region and resulting in an intermediate FRET efficiency.

In this weakened, partially engaged state, the allosteric input from PIP_2_ binding in the C-lobe is sufficient to promote complete dissociation of the C-tail to the active, low-FRET conformation. This partially engaged state also reduces the binding energy for a GPCR tail, as evidenced by an approximately 30-fold increase in sensitivity to V2Rpp observed for 3A βarr1 by smFRET (EC_50_: ∼400 nM versus 11.6 μM; Figure S6A), and the lower specificity for phosphorylated GPCRs^50^.

For the middle C-tail mutation, we chose to examine the effect of mutating R393, a residue that forms part of the so-called polar core of arrestin^51^. The polar core helps to lock the N- and C-terminal domains in the inactive orientation in the absence of a GPCR and is composed of electrostatic interactions between residues D26, R169, D290, and D297 in the N- and C-domains, and residue R393 in the middle C-tail^52,53^ (Figure 5A). Disruption of this charge balance during activation enables interdomain twisting and helps to dissociate the βarr1 C-tail. R393Q was found to have a relatively weak phenotype, showing only a modest increase in binding to non-phosphorylated activated GPCRs^51^. Compared to the 3A mutation, R393Q caused only a small increase in mid-FRET occupancy relative to the parent construct, with the majority of the βarr1 population remaining in the high-FRET, inactive state (Figures 5C and S6C). Consistent with this observation, R393Q βarr1 also showed less enhancement in V2Rpp binding affinity compared to 3A in smFRET measurements (Figure S6C). However, it could still be fully activated by V2Rpp. These data suggest that while R393Q disrupts anchoring of the C-tail in the N-domain, it does so to a lesser extent than the 3-EI mutation.

The addition of 100 μM PIP_2_ to R393Q βarr1 led to a nearly complete loss of high-FRET occupancy, along with a substantial increase in the low-FRET state (Figures 5C and S6C). These changes indicate a similar relative increase in C-tail displacement compared to the 3A mutant, with a change in fractional low-FRET of 0.3 and 0.33, respectively. Similarly, FP binding measurements showed that R393Q, like 3A, increased the affinity of PIP_2_ for βarr1, albeit to a lesser extent (Figure 5D). Inspection of individual trajectories in the presence of PIP_2_ revealed transitions between all three states, though transitions primarily occurred between the high- and the mid-FRET states, as well as the mid- and the low-FRET states (Figure 5C). These data suggest that PIP_2_ binding cannot fully disengage the C-tail directly from the inactive, high-FRET state, even in the context of activated mutants of βarr1. Based on our data, we propose that PIP_2_ binding in the C-lobe shifts the βarr1 C-tail to a partially activated state of disengagement. Full C-tail disengagement appears to require βarr1 to already be in the mid-FRET state for PIP_2_ binding to promote a transition to low-FRET. This supports a model of sequential transitions through the C-tail intermediate, consistent with the transition density plot of R393Q imaged in the presence of 100 μM PIP_2_ (Figure 5C).

### C-tail activation enhances PIP binding affinity

Having demonstrated that PIP_2_ binding allosterically influences the conformation and dynamics of the C-tail, we reasoned that C-tail mutations may also alter PIP_2_ binding to βarr1. Using FP to measure PIP_2_ binding, we found that mutations that eliminate or destabilize the auto-inhibited state of βarr1 either by removing the C-tail entirely (ΔCT) or compromising either of the anchor points (R393Q, 3A) increased the affinity of PIP_2_ for βarr1 by an order of magnitude (Figure 5D). These data demonstrate a linked allosteric network between the concave surface of the C-lobe and the N-domain. They also suggest that PIP_2_ preferentially binds to an active-like conformation of βarr1, with either a disengaged or partly disengaged C-tail. Supporting this model, the low-FRET dwell times representing the PIP_2_-bound state of the 3A mutant were significantly prolonged compared to the analogous mid-FRET dwells of the parent sensor (ca. 11 s versus 2 s; each at 100 μM PIP_2_), indicating that the 3A mutation likely enables PIP_2_ to remain bound in the C-lobe for a longer duration.

### Global βarr1 conformational changes induced by PIP_2_

The data from FP and smFRET support an allosteric connection between the PIP_2_ binding site in the C-lobe and the βarr1 C-tail; however, questions about the broader structural changes within βarr1 that occur upon PIP_2_ binding remained. To assess this, we used hydrogen-deuterium exchange mass spectrometry (HDX-MS) and compared the effects of PIP_2_ binding on the parent (henceforth WT), 3A and R393Q βarr1 constructs. We obtained >90% sequence coverage in our MS experiments for all the tested constructs, with many positions showing coverage by multiple overlapping peptides, allowing for improved resolution of structural information. To visualize the changes in exchange properties, we simulated deuterium exchange kinetics^54^ and computed protection factors (PF) using a Monte Carlo approach to obtain a best-fit residue-level representation of global conformational changes in βarr1 between conditions (see Methods). While we had limited peptide coverage of the N-terminal β-strand and the C-terminus, around the locations of our smFRET probes, our data recapitulated and extended prior findings.^55^ In comparing βarr1 WT and 3A, we observed an expected increase in dynamics across βarr1 resulting from the disruption of the three-element anchor, with major changes occurring in the gate, finger and middle loops; however, unexpectedly, we also saw a large increase in exchange in the C-lobe β-strands that comprise the PIP_2_ binding site, an effect also seen for the WT/R393Q dataset. Second, while R393Q showed many similar increases in dynamics, albeit to a lesser extent than 3A, there were also differences, such as at the N-terminal region of the N-lobe and the 160 loop^56^ in the central crest. These data are suggestive that R393Q and 3A exhibit different conformational ensembles in their basal state, consistent with our smFRET experiments.

The addition of PIP_2_ to WT βarr1 led to a global decrease in deuterium uptake. This was most pronounced along the concave surface of the C-lobe, where binding of the phosphoinositide headgroup was expected to reduce solvent accessibility (Figures 6A and 6B). Beyond the C-lobe, we observed reduced dynamics in regions of the central crest^47^, including the finger loop, which is expected to be more ordered upon βarr1 activation. We also observed a slight reduction in dynamics across the N-domain, which we reason is a consequence of partial interdomain twisting^4^ and the absence of canonical C-tail release. Consistent with this, a recent HDX-MS study on βarr2 also showed PIP_2_ binding decreased exchange in the extended N-domain groove around the distal C-tail^34^.

**Figure 6.**
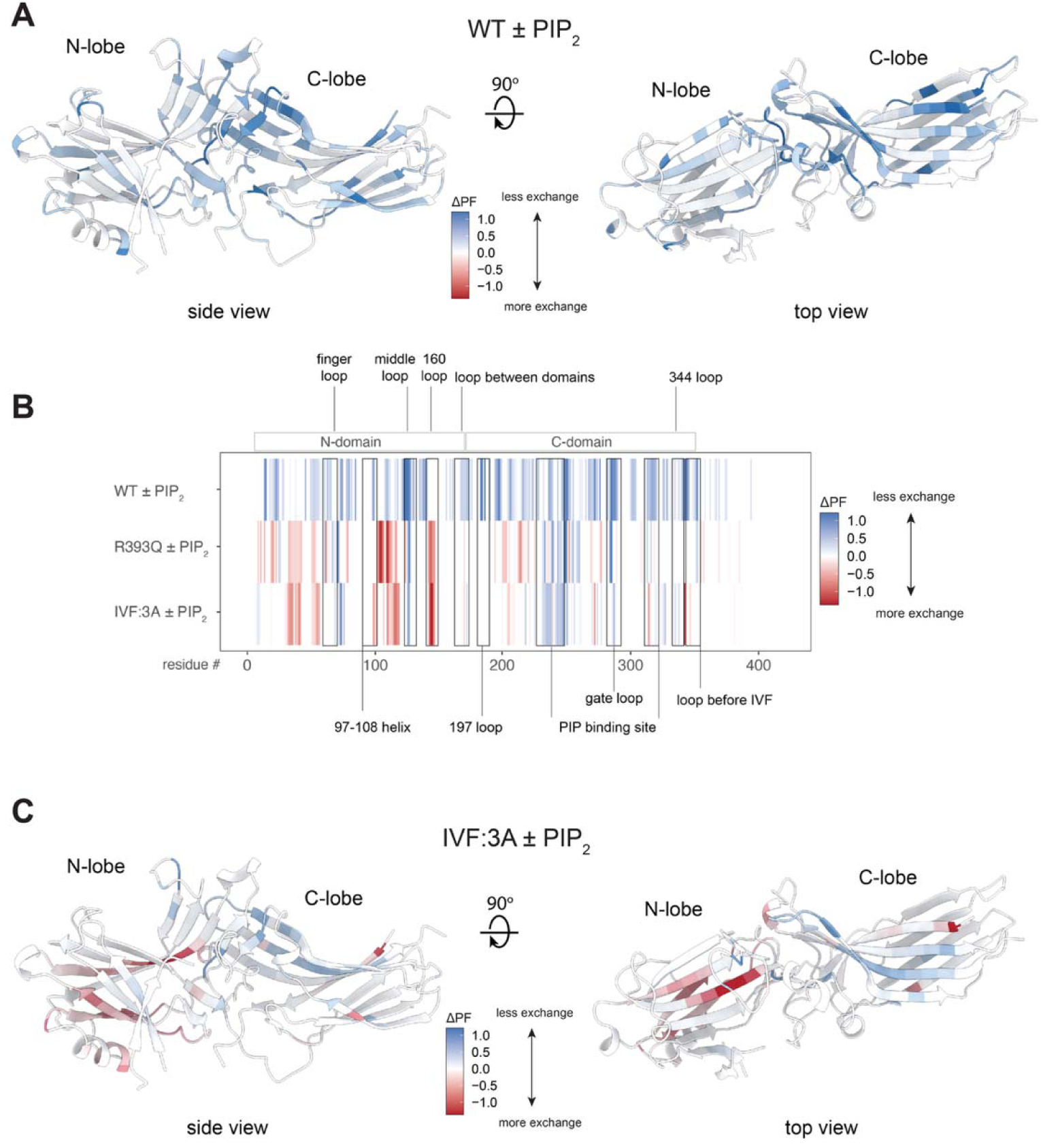
HDX-MS shows conformational changes in βarr1 induced by PIP2 binding. HDX-MS measurements of βarr1 constructs were taken at various time points. The resulting deuterium uptake profiles were modeled, yielding a protection factor at each residue (STAR Methods). (A and B) Residues showing significant changes in deuterium exchange mapped onto the inactive state structure of βarr1 (PDB: 1G4M) for (A) WT and (B) 3A constructs. Scale bar, normalized protection factor (PF) change in response to PIP2: ΔPF = PFbasal - PFPIP2. (C) Linear map (residue by residue) of ΔPF upon addition of PIP2 for various βarr1 constructs. See also Figure S7.

In contrast to WT βarr1, the R393Q and 3A constructs showed similar deuterium exchange profiles (Figures 6B, 6C, S7A, and S7B) in the presence of PIP_2_. While we observed a reduction in deuterium uptake on the concave surface of the C-lobe for both, like WT βarr1, the concave surface of the N-lobe showed a substantial increase in deuterium content. This is consistent with our smFRET experiments where PIP_2_ addition to both R393Q and 3A βarr1 resulted in an increased population of low-FRET, C-tail-dissociated state (Figures 5B and 5C). We note that the significant difference in HDX profiles between the WT + PIP_2_ condition and basal 3A mutant conditions (cf. Figure S7B and Figure 6B) suggests that while both conditions promote movement of the C-tail, the underlying mechanisms driving this process differ, and PIP_2_ does not simply act to disrupt the 3EI.

To better understand movements in the middle region of the βarr1 C-tail, we used an additional smFRET labeling site (G398C) located in that region (Figure S8A). Single-molecule experiments using this construct (βarr1-12-398-AF488-AT647) (Figure S8B and S8C) showed that under basal conditions, the C-tail predominantly adopts a single broadly defined mid-FRET state (∼0.4). Treatment with an EC_50_ concentration of V2Rpp (0.75 μM) partially shifted the FRET distribution to a low-FRET (∼0.2) state, as expected for activation. Interestingly, treatment with 100 μM PIP_2_ promoted transitions from the mid- to a high-FRET (∼0.6) state, with near-total (>80%) occupancy of the high-FRET state at the population level (Figure S8B and S8C). This increase in FRET efficiency suggests that PIP_2_ binding repositions the middle C-tail down within the N-domain (Figure S8D), which would be consistent with the observed effects with our 12-387 sensor and supported by our HDX-MS data. Notably, movement of the C-tail within the N-domain groove has been observed in MD simulations^45^. If the proximal C-tail is partly released, resulting in a mid-FRET state, the C-tail presumably can remain anchored via its middle segment that contacts the polar core.

### Role of the intermediate state in βarr1 activation

Given the effect of PIP_2_ on the conformation of βarr1, and its ability to shift βarr1 towards a more active-like conformation, we aimed to understand the significance of this active-like conformation in cells. Using a previously established split luciferase complementation assay,^25,57^ we measured βarr1 recruitment to several GPCRs, including those that require βarr1 co-binding of PIP_2_ for recruitment and those where this is dispensable (Figure 7A)^25^. Specifically, we compared the relative recruitment of WT βarr1 to 3Q βarr1, the variant incapable of binding to PIP_2_ both in the context of the native βarr1 C-tail or with the 3A proximal C-tail mutations. We measured the time-dependent recruitment of these βarr1 construct pairs and generated corresponding concentration-response curves for 5 GPCRs (Figure 7B), two class A (weak βarr-interactors) GPCRs, two class B GPCRs (strong βarr-interactors) and a chimeric GPCR. The two class A GPCRs, β1AR and NTSR1-6A^25^, a variant of NTSR1 lacking C-terminal phosphosites, showed pronounced loss of βarr1 recruitment for 3Q βarr1 while the two class B GPCRs, NTSR1 and V2R, showed little or no difference in 3Q βarr1 recruitment. As seen previously, the chimeric β2AR-V2C receptor showed an intermediate effect between the two^25^. Introduction of the proximal C-tail 3A mutations fully rescued the loss of recruitment of 3Q βarr1 to β1AR, NTSR1-6A and β2AR-V2C (Figure 7C).

**Figure 7.**
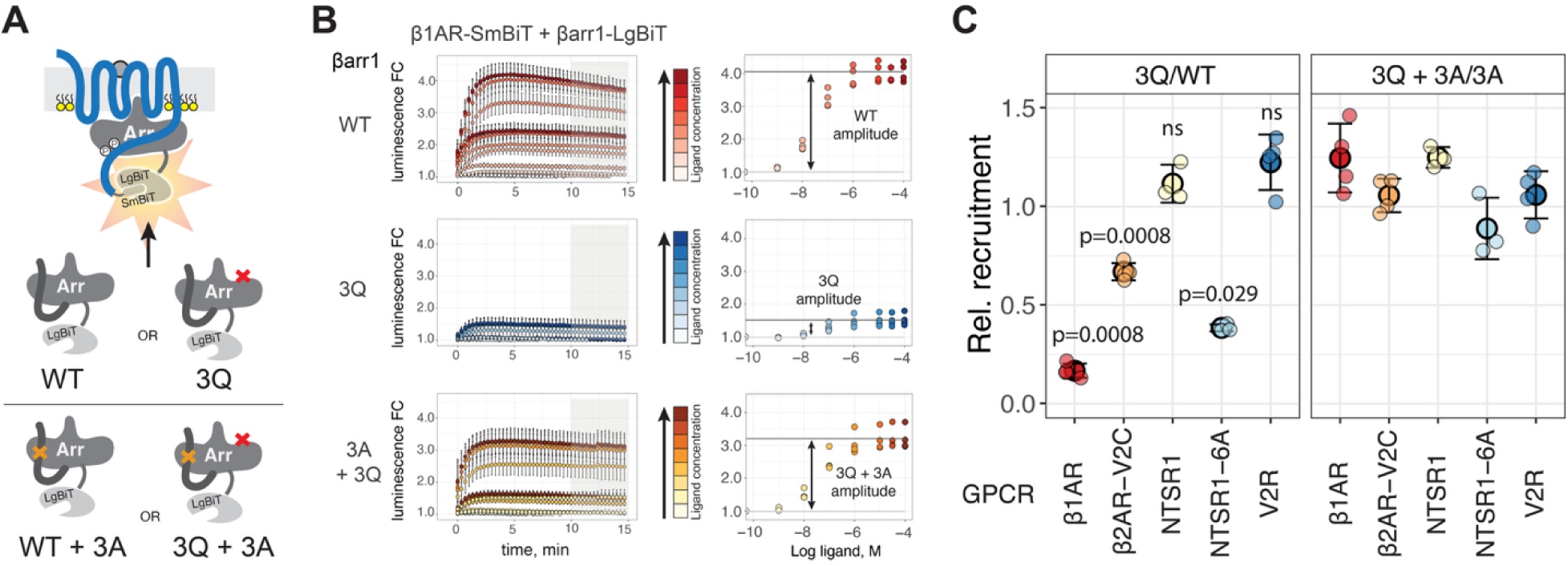
C-tail mutations rescue βarr1 recruitment to class A GPCRs when PIP_2_ binding is eliminated. (A) Cartoon showing split luciferase complementation assay to measure βarr1 recruitment to GPCRs in live cells. Red X denotes K232/R236/K250 to 3Q triple mutant. Orange X denotes I377A/V378A/F379A triple mutant. (B and C) (B) Time-dependent recruitment response for different βarr1 constructs as a function of ligand concentration and (C) the relative recruitment to various GPCRs of PIP-binding deficient mutants of WT (left) and 3A (right) βarr1. Error bars in B (left) and C are s.d., n = 4-5 replicates. p values correspond to unpaired t-tests with Welch’s correction between 3Q/WT and (3Q+3A)/3A conditions for each GPCR. See also Figure S9.

These findings demonstrate that compromising the βarr1 C-tail is sufficient to overcome the requirement for membrane phosphoinositides for βarr1 recruitment to class A GPCRs, which we attribute to increasing the strength of the GPCR-core interactions with βarr1 when phosphorylation is limiting. This could be manifest in either of two ways: the 3A mutation may increase PIP_2_ binding affinity to βarr1 3Q (Figure 5D), or the conformational changes conferred by the 3A mutation can increase probability of productive GPCR interactions, or some combination of these mechanisms. Our in vitro experiments reveal that PIP_2_ and the 3A mutation result in similar βarr1 C-tail conformations with predominant occupancy of the intermediate FRET state (Figures 4 and 5, respectively). In this context, we speculate that PIP_2_ may be necessary to enhance recruitment to weakly βarr-interacting receptors by triggering conformational changes, including partially disengaging the C-tail. This mechanism may also be important for assembly of the core- and tail-bound, fully assembled state of a GPCR-βarr1 complex^25^.

To further explore how the intermediate state might facilitate βarr1 engagement of GPCRs, we performed surface plasmon resonance (SPR) measurements probing Fab30 binding to βarr1 3A. Previous work showed that apo WT βarr1 bound weakly to Fab30, with binding modestly enhanced by PIP_2_, and strongly enhanced by V2Rpp^25^. In contrast, we found that binding of Fab30 to βarr1 3A showed little or no enhancement with addition of V2Rpp (Figures S9A and S9B). Structural analysis suggests that optimal Fab30 binding requires proper positioning of the peptide chain in βarr1 N-domain, supported by the fact that Fab30 binding to βarr1 ΔCT without V2Rpp was significantly less enhanced than to 3A (Figures S9C). These data suggest that sliding of the C-tail in a manner that produces a mid-FRET state (via PIP_2_ or the 3A mutation) reflects a more activated conformation of βarr1. Additionally, smFRET measurements showed that, in the absence of phosphopeptide, Fab30 binds to and stabilizes the mid-FRET conformation (Figure S9D), further supporting the idea that this βarr1 intermediate is recognized as active by the Fab.

Together, these data show that membrane phosphoinositides modulate βarr1’s conformational ensemble to promote an active-like state necessary for core-engagement of a GPCR, especially in the context of weakly phosphorylated receptors whose C-termini cannot independently displace the βarr1 tail from its inactive position.

## DISCUSSION

In this study, smFRET imaging of βarr1 reveals that the C-tail possesses intrinsic basal dynamics and samples at least three distinct states^14^, rather than the previously described two-state system^5^. Importantly, these intermediately active states can be potentiated by allosteric modulators, such as PIP_2_, and are structurally distinct from the canonical active or inactive states observed structurally^43,58–60^. Despite >250 arrestin structures, only one recent report^61^ shows almost the entire C-tail of Arr1 bound within the N-lobe. However, even these structures fail to explain how short-lived and metastable conformational states of βarrs may result from movement or partial C-tail displacement. The data presented herein offer a mechanistic basis for understanding the nature of discrete βarr1 conformations that exist between the well-characterized extrema, and which may better explain how membrane inputs^25,38^, phosphorylation patterns^4,62^, and differences in receptor binding poses^10^ may synergize to produce multiple distinct βarr states capable of eliciting specific downstream responses^4,6,7^.

Though the proximal βarr1 C-tail primarily adopts an inactive, autoinhibited conformation, it is basally dynamic with rapid excursions to partially and fully disengaged active states (Figures 2A-E). We believe these transitions explain how autoinhibition is briefly attenuated, allowing for initial engagement with an active, phosphorylated receptor to occur^56^. However, given the range of measured phosphopeptide affinities, it is likely that most class A typology (weakly βarr-interacting) GPCRs^25,37,63,64^ would lack suitable C-terminal phosphorylation to effectively capture βarr1 during brief, active-state fluctuations amid excursions to the plasma membrane^32^. This is consistent with GPCRs of this typology requiring additional activating inputs from the plasma membrane, such as PIP_2_, to achieve effective interactions^25^. In addition to the conformational priming, binding of βarr1 to membrane regions rich in PIP_2_ may extend the lifetime of βarrs at the membrane^35,36^, increasing the likelihood of productive collisions with an agonist-bound GPCR^32^.

While allosteric communication within βarrs has been suggested^65^, our data provide clear evidence for a strong connection between the C-lobe of βarr1 and its C-tail. We demonstrate that PIP_2_ binding in the C-lobe allosterically destabilizes the fully inhibited state of the βarr1 C-tail and promotes transition to an intermediate state (Figures 4B-E). This state likely follows from the dissociation of the C-tail’s proximal segment surrounding the 3-EI, as mutations in this region induced a state with similar FRET efficiency (Figures 5A and 5B), and rescued recruitment of PIP-binding-deficient mutants in cells (Figure 7C). Additionally, we observed that a V2Rpp derivative, P_56_, which contains the phosphorylation sites necessary to bind βarr1 and displace the C-tail only in its proximal region, similarly shifted the βarr1 C-tail to the intermediate state. This further supports our model that the intermediate state involves proximal, but not distal, C-tail release and provides new evidence for non-switch-like behavior for the βarr1 C-tail. Our data indicates that this partial activation involves just over half the βarr1 C-tail displacement observed in the fully active state, consistent with earlier double electron-electron resonance (DEER) spectroscopy measurements^42^. Notably, MD simulations have suggested that C-tail movement, but not release, may expose phosphate-binding sites crucial for facilitating initial engagement of the Rpp tail^4,62^.

Using C-tail displacement as a proxy for βarr1 activation, our data suggests that binding of different peptide and lipid ligands (i.e., V2Rpp, PIP_2_ and P_56_), or the presence of mutations (i.e., R393Q and 3A), can result in varying degrees of activation (Figures 3 and 4). These data support a cooperative model for crosstalk between the PIP_2_ and GPCR phosphorylation-mediated binding, where dissociation of the βarr1 C-tail, induced by phosphopeptides or mutations, is differentially enhanced by PIP_2_ binding, potentially leading to the adoption of the fully active state that may not be inaccessible with either input alone (Figures S5 and 5A-C). Similar observations have been made for βarr2^34^. In this way, phosphorylation pattern-dependent regulation of βarr activity could be modulated by spatial variations in the local concentration of PIP_2_ or other membrane phospholipid pools^66,67^. Additional interactions with the plasma membrane^32,33,67^, inositol hexaphosphate (IP_6_)^68^, and the native glycosaminoglycan heparin^45^ may further influence the activation and conformation of βarrs in cells.

Although a prior smFRET study reported both a lack of basal dynamics and no effect of PIP_2_ on the βarr1 C-tail^45^, their system employed different labeling sites (176-397) and dyes (LD550/LD650, analogous to Cy3/5) than the ones used in this work. Furthermore, our findings that PIP_2_ alters the βarr1 C-tail by smFRET are corroborated by evidence from us and other groups, which show that PIP_2_ has an effect on both βarr1 and βarr2 conformation using NMR^33^, HDX-MS^34^, and ensemble fluorescence^25,35^.

Our findings suggest that βarr1 can integrate multiple activating inputs, enabling fine-tuning of its C-tail conformation, precisely gating interactions with a target GPCR. This conformational flexibility sheds light on how just two βarrs can engage the over 800 human GPCRs. Moreover, these distinct βarr1 conformational states may be the source of differences that are recognized by downstream effector proteins, including clathrin and adaptin proteins^69^, various kinases^70,71^ – many of which also interact with PIP_2_^72–74^, and other arrestin interactors^75^. We believe future studies will further refine these hypotheses and delineate how diverse pathways are integrated by βarrs at the level of its structural dynamics to direct signaling towards >100 downstream effectors.

## Limitations of this study

Dynamics studies are limited to in vitro experiments that use a soluble derivative of membrane PI(4,5)P_2_ (diC8 versus native alkyl chains). These experiments required high micromolar concentrations of PIP_2_, which may more potently affect βarr1’s conformation at the plasma membrane. However, we note that the concentration of PIP_2_ in mammalian cells (if it were dissolved in the cytosol) is high, approximately 10 μM, and it is significantly higher in nanodomains^76,77^ around some GPCRs^78,79^. In addition, smFRET is a precise measure of relative distance changes but not a good measure of absolute distance^80^. Therefore, further studies are needed to characterize the structure of the βarr1 intermediate state with high spatial resolution. Future studies investigating the βarr1 C-tail at sub-10 ms time scales could expand our understanding of its dynamics. Finally, although our cell-based recruitment assays offer important mechanistic insights, they were conducted in HEK293 cells with overexpression of the protein components.

## ACKNOWLEDGMENTS

We thank Dr. Elizabeth White for technical laboratory assistance; Dr. Shoji Maeda (Stanford University) for providing purified Fab30 protein; D. Hilger for providing BirA enzyme; Kayo Sato, Shigeko Nakano, and Ayumi Inoue (Tohoku University) for their assistance with plasmid construction and cell-based GPCR assays. This work was supported in part by National Institutes of Health grants K99GM147609 (to J.J.), R01GM143554 (to S.C.), and R01NS028471 (B.K.K.). Additional support to B.K.K. was provided by the Mathers Foundation. Additional support to S.C. was provided by the Eleftheria Foundation. B.K.K. is a Chan-Zuckerberg Biohub Investigator. J.J. is a Damon Runyon Fellow supported by the Damon Runyon Cancer Research Foundation (DRG-2318-18). R.V.S was an Interdisciplinary Scholar of the Wu-Tsai Neuroscience Institute (Stanford University). M.M. was supported by an American Heart Association postdoctoral fellowship (17POST33410958). A.I. was funded by KAKENHI JP21H04791, JP24K21281 and JP25H01016 from the Japan Society for the Promotion of Science (JSPS); JP22ama121038 and JP22zf0127007 from the Japan Agency for Medical Research and Development (AMED); JPMJFR215T and JPMJMS2023 from the Japan Science and Technology Agency (JST); The Uehara Memorial Foundation.

## AUTHOR CONTRIBUTIONS

Conceptualization, J.J., B.K.K. and R.V.S.; Methodology, J.J., J.D., R.V.S.; Code, J.J., J.D., R.V.S.; Formal Analysis, J.J., J.D., Y.S., R.V.S.; Investigation, J.J., J.D., Y.S., A.I., D.H.S., W.L., R.V.S.; Resources, M.M., R.V.S., Data Curation, J.J., J.D., R.V.S.; Writing – Original Draft, J.J., J.D., R.V.S.; Writing – Review & Editing, J.J., J.D., Y.S., A.I., D.H.S., W.L., M.M., R.V., S.C., B.K.K., R.V.S.; Visualization, J.J., J.D.; Supervision, R.V., S.C., B.K.K., R.V.S.; Funding Acquisition S.C., B.K.K.

## METHODS

### General

The V2Rpp (ARGRpTPPpSLGPQDEpSCpTpTApSpSpSLAKDTSS) and P_56_ (ARGRTPPSLGPQDESCTpTApSpSpSLAKDTSS) peptides were obtained by custom peptide synthesis (Tufts University Core Facility). The concentration of V2Rpp stocks was determined by reaction with Ellman’s reagent as previously described^4^. Fab30 was expressed and purified as previously described^43^. Soluble PIP derivatives were purchased from Avanti Polar Lipids as powders and reconstituted in 50 mM HEPES pH 7.4 to a stock concentration of 1-5 mM. For mass spectrometry, LC-MS grade water, LC-MS 0.1% formic, and LC-MS grade acetonitrile with 0.1% formic acid in water were purchased from Fisher Scientific (Hampton, NH). Guanidine hydrochloride and citric acid were purchased from Sigma-Aldrich (St. Louis, MO). Deuterium oxide (99+ %D) was purchased from Cambridge Isotope Laboratories (Tewksbury, MA).

### Plasmid Construction

For cell-based assays, we used human, full-length GPCR plasmids cloned into the pCAGGS vector or the pcDNA3.1 vector derived from a previous study^81^. NTSR1 constructs were N-terminally FLAG epitope-tagged with a linker (M<u>DYKDDDDK</u>GTELGS; the FLAG epitope tag is underlined) and inserted into the pcDNA3.1 vector. β1AR, β2AR-V2C and V2R constructs had an N-terminal FLAG epitope tag with a preceding HA-derived signal sequence and a flexible linker (MKTIIALSYIFCLVFA<u>DYKDDDDK</u>GGSGGGGSGGSSSGGG) and inserted into the pCAGGS vector. For the direct NanoBiT-based βarr recruitment assay, human full-length βarr1 was N-terminally LgBiT-fused with the same flexible linker and inserted into the pCAGGS vector (LgBiT-βarr1). GPCRs were C-terminally SmBiT-fused with the flexible linker (GGSGGGGSGGSSSGG<u>VTGYRLFEEIL</u>; the SmBiT is underlined) and inserted into the pCAGGS vector (GPCR-SmBiT).

### NanoBiT-β-arrestin recruitment assays

Direct βarr1 recruitment was measured as follows. HEK293A cells (Thermo Fisher Scientific) were seeded in a 6-cm culture dish (Greiner Bio-One) at a concentration of 2 x 10^5^ cells per ml (4 ml per dish hereafter) in DMEM (Nissui Pharmaceutical) supplemented with 10% FBS (Gibco), glutamine, penicillin, and streptomycin, one day before transfection. The transfection solution was prepared by combining 5 µl of polyethylenimine (PEI) solution (1 mg/ml) and a plasmid mixture consisting of LgBiT-βarr1 variant (500 ng) and C-terminally fused-SmBiT GPCR (500 ng) constructs in 200 µl of Opti-MEM (Thermo Fisher Scientific). After 24 hours the transfected cells were harvested with 0.5 mM EDTA-containing Dulbecco’s PBS, centrifuged, and suspended in 2 ml of Hank’s balanced saline solution (HBSS) containing 0.01% bovine serum albumin (BSA fatty acid-free grade, SERVA) and 5 mM HEPES (pH 7.4) (assay buffer). The cell suspension was dispensed in a white 96-well plate (Greiner Bio-One) at a volume of 80 ml per well and loaded with 20 µl of 50 µM coelenterazine (Carbosynth), diluted in the assay buffer. After a 2 h incubation at room temperature, baseline luminescence was read using a SpectraMax L, 2PMT model (Molecular Devices). Following this, 20 ml of 6x ligand serially diluted in the assay buffer was manually added. The following ligands were used: isoproterenol (iso) for β1AR and β2AR-V2C, Arginine vasopressin for V2R, and neurotensin for NTSR1 and NTSR1-6A. The plate was immediately read for the second measurement as a kinetics mode and luminescence counts recorded for 15 min with an accumulation time of 0.18 sec per read and an interval of 20 sec per round. For every well, the recorded kinetics data were first normalized to the initially recorded baseline luminescence signal.

### Analysis of cell-based recruitment data

NanoBiT data were analyzed by converting kinetic data into concentration-response data by determining an average fold-change (relative to signal pre-stimulation) from 10-15 minutes post-agonist addition. At least three independent experiments were performed for each GPCR-arrestin sensor combination. Concentration-dependent data from two technical replicates for each independent experiment were collectively fit to a four-parameter log logistic function (LL2.4) provided in the drc package (v 3.0-1)^82^ of the statistical environment R^83^. This equation, of the form: 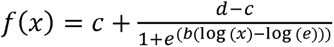 provides pre- and post-transition values, c and d, respectively, that define the amplitude response for that assay. Uncertainty is propagated from the error of each amplitude given by RSS of the top and bottom of the fits.

### β-arrestin expression and purification

The parent construct for β-arrestin 1 (βarr1) is the long splice variant of human, where all cysteine residues are removed by mutation (C59V, C125S, C140L, C150V, C242V, C251V, C269S). Additional mutations were added by site-directed mutagenesis and these proteins were expressed and purified as the parent protein. This construct is modified with an N-terminal 6x Histidine tag, followed by a 3C protease site, a GG linker, AviTag and GGSGGS linker. The sequence was codon-optimized for expression in *E. coli* and cloned into a pET-15b vector. Point mutations were prepared using site-directed mutagenesis. βarr1 (ΔCT) was prepared by truncating βarr1 after residue 382. All βarr1 constructs used were prepared as follows: NiCo21(DE3) competent *E. coli* (NEB) were transformed, and large-scale cultures were grown in TB + ampicillin at 37°C until an OD_600_ of 1.0. Cells were then transferred to room temperature and induced with 25 µM IPTG when the OD_600_ reached 2.0. Cells were harvested 20 h post induction and resuspended in lysis buffer [50 mM HEPES pH 7.4, 500 mM NaCl, 15% glycerol, 7.13 mM 2-mercaptoethanol (BME)] to a final volume of 40 mL/L of cells. Cells were lysed by sonication and the clarified lysate applied to nickel sepharose and batch incubated for 1.5h at 4°C. The resin was washed with 10 column volumes of wash buffer (20 mM HEPES pH 7.4, 500 mM NaCl, 10% glycerol, 7.13 mM BME) + 20 mM imidazole, followed by 10 column volumes of wash buffer + 40 mM imidazole. The protein was then eluted with 5 column volumes of wash buffer + 200mM imidazole and dialyzed overnight in 100x volume of dialysis buffer (20 mM HEPES 7.4, 200 mM NaCl, 2 mM BME, 10% glycerol) in the presence of 1:10 (w:w) of 3C protease. The digested protein was then subjected to reverse-Nickel purification and diluted with a dialysis buffer containing no NaCl to bring the NaCl concentration to 75mM. The protein was then purified by ion exchange chromatography (mono Q 10/100 GL, GE Healthcare), followed by SEC using a Superdex 200 increase 10/300 GL column (GE Healthcare) with SEC buffer (20 mM HEPES pH 7.4, 300 mM NaCl, 10% glycerol). Purified protein was concentrated to between 100-300 mM using a 30 kDa MWCO spin concentrator and aliquots were flash-frozen in liquid nitrogen and stored at -80°C until use.

### β-arrestin labeling and biotinylation

Following SEC, elution peak fractions were pooled to a concentration of 10-20 µM and labeled with a 1:3 mixture of AlexaFluor488-C5 maleimide and ATTO647N maleimide, respectively. Fluorophores were dissolved in DMSO to 25-40 mM and added at 10x molar excess over protein, then allowed to react for 1 h at room temperature prior to quenching with L-Cysteine (10x molar excess over fluorophore). The labeling reaction was further incubated for 10 minutes after cysteine addition, after which samples were spin filtered and subjected to a second round of size-exclusion chromatography, as detailed above, to remove free dye. The purified protein was concentrated to between 100-300 µM using a 30 kDa spin concentrator, and aliquots were flash-frozen in liquid nitrogen and stored at -80°C until use.

Arrestins (SEC-pure) were biotinylated using recombinant BirA enzyme, according to commercial protocols (Avidity), with exception that biotinylation was carried out for 12 h at 4°C, rather than 30°C. After biotinylation was complete, the reaction was flowed over 100 mL (packed) of nickel Sepharose, equilibrated in arrestin SEC buffer and supplemented with 10 mM imidazole, then washed with 200 mL of the equilibration buffer. The combined flow-through and wash fractions were then purified by size-exclusion chromatography as described above.

### Fab30 expression and purification

Fab30^43^ was cloned into pFastBac-dual with an octa-histidine tag added to the C-terminus of the heavy-chain subunit (with an intervening AAA linker) and a GP67 secretion signal added to the N-terminus of both the heavy- and the light-chains. Fab30 was expressed by Hi5 insect cells (Expression Systems) into the media using a FastBac-derived baculovirus. Cells were infected at a density of 3x10^6^ cells/mL and harvested 72 hrs post-infection. Cells were pelleted by centrifugation, and the supernatant was transferred to a large beaker with constant stirring. Tris pH 7.5 was added to a final concentration of 50 mM, followed by NiCl_2_ (to 1 mM) and CaCl_2_ (to 5 mM). A heavy precipitate will form. Add protease inhibitors and stir at room temperature for 45 minutes. Any precipitate was removed by centrifugation, and the clarified supernatant was applied to nickel sepharose 2 mL resin/L of supernatant. The resin was washed with 100 mL of 20 mM HEPES pH 7.5, 500 mM NaCl, 20 mM imidazole, then with 100 mL of 20 mM HEPES pH 7.5, 100 mM NaCl, 20 mM imidazole. Fab30 was eluted with 20 mM HEPES pH 7.5, 100 mM NaCl, 250 mM imidazole. Pooled fractions containing Fab30 were concentrated and subjected to polishing by SEC on a Superdex 200 pg 16/600 GL column (GE Healthcare) in 20 mM HEPES, pH 7.5, 100 mM NaCl. Peak fractions were pooled and concentrated to 200-300 mM, supplemented with glycerol to 15% v/v final, and aliquots were flash-frozen and stored at - 80°C until use.

### NTSR1 expression and purification

Full-length human NTSR1 was modified with an N-terminal Flag tag followed by an octa-histidine tag and cloned into pFastBac1 vector. NTSR1 was expressed in Sf9 insect cells (Expression Systems) using a FastBac-derived baculovirus. Cells were infected at a density of 4x10^6^ cells/mL and harvested 60 hrs post-infection. Cells were lysed in hypotonic buffer (10 mM HEPES, pH 7.4, and protease inhibitors) and solubilized at 4°C for 2 hours in a buffer containing 1% lauryl maltose neopentyl glycol (LMNG, Anatrace), 0.1% cholesteryl hemisuccinate tris salt (CHS, Steraloids), 0.3% sodium cholate (Sigma), 20 mM HEPES 7.4, 500 mM NaCl, 25% glycerol, iodoacetamide (to cap cysteine residues) and protease inhibitors. Insoluble debris was removed by centrifugation and the supernatant was incubated with Ni-NTA (Qiagen) resin for 1 hour at 4 °C. The resin was washed in batch with buffer containing 0.01% LMNG, 0.001% CHS, 0.003% sodium cholate, 20 mM HEPES pH 7.4, 500 mM NaCl, 10 mM imidazole and eluted with the same buffer supplemented with 200 mM imidazole, 2 mM CaCl_2_ and 10 μM NTS_8-13_ (Acetate salt, Sigma). The eluate was loaded onto M1 FLAG immunoaffinity resin and washed with buffer containing 0.01% LMNG, 0.001% CHS, 0.003% sodium cholate, 20 mM HEPES pH 7.4, 500 mM NaCl, 10 mM imidazole, 0.1 μM NTS_8-13_ and 2 mM CaCl_2_. The receptor was eluted with buffer containing 100 mM NaCl, 20 mM HEPES pH 7.4, 0.005% LMNG, 0.005% CHS, 1 μM NTS_8-13_, 0.2 mg/mL flag peptide (DYKDDDDK) and 5 mM EDTA. Elution fractions containing receptor were pooled and subjected to polishing by SEC on a Superdex 200 Increase 10/300 GL column (GE Healthcare) in 20 mM HEPES, pH 7.4, 100 mM NaCl, 0.0025% LMNG, 0.00025% CHS, and 0.1 μM NTS_8-13_. Peak fractions were pooled and concentrated to 200 μM, and aliquots were flash-frozen and stored at -80°C until use.

### GRK5 expression and purification

Full length human GRK5 was modified with a C-terminal hexa-histidine tag and cloned into pVL1392 vector for baculovirus production. GRK5 was expressed and purified as previously described^25^. Briefly, Sf9 insect cells (Expression Systems) were infected with a BestBac-derived baculovirus at a density of 3.5 x 10^6^ cells/mL and harvested 48 hours post infection. Cells were resuspended, lysed by sonication and the supernatant was applied to Ni-NTA resin. The resin was washed with lysis buffer and GRK5 eluted with lysis buffer supplemented with 200 mM imidazole. The combined eluate was then subjected to cation-exchange chromatography using a MonoS 10/100 column (GE healthcare) and eluted with a linear gradient of NaCl. Fractions containing GRK5 were combined and run on a Superdex 200 10/300 GL column (GE healthcare). GRK5 was aliquoted, flash frozen, and stored at -80 °C until use.

### NTSR1 in vitro phosphorylation

NTSR1 (2.5 μM) was equilibrated in phosphorylation buffer (20 mM bis-tris propane (BTP) pH 7.5, 35 mM NaCl, 5 mM MgCl2, 20 µM NTS_8-13_, 20 μM 08:0 PI(4,5)P2, 0.05 mM TCEP, 0.002% MNG, 0.0002% CHS) at 25 °C with gentle mixing for 1 h. GRK5 was added to the reaction to a final concentration of 200 nM, and briefly incubated while the reaction was warmed from 25 °C to 30 °C. ATP was added to a final concentration of 1 mM. Upon completion, the reaction was supplemented with CaCl_2_ to a final concentration of 2 mM and applied to an equilibrated M1 FLAG immunoaffinity resin and washed with buffer containing 0.004% LMNG, 0.004% CHS, 20 mM HEPES pH 7.4, 100 mM NaCl, 0.2 μM NTS_8-13_, 2 mM CaCl_2_. The receptor was eluted with buffer containing 100 mM NaCl, 20 mM HEPES pH 7.4, 0.004% LMNG, 0.004% CHS, 0.2 μM NTS_8-13_, 0.2 mg/mL Flag peptide, 5 mM EDTA), followed by SEC using a Superdex 200 increase 10/300 GL column (GE Healthcare) with SEC buffer (20 mM HEPES pH 7.4, 100 mM NaCl, 0.004% LMNG, 0.0004% CHS).

### Analytical fluorescence-detection size-exclusion chromatography

In a final volume of 20 mL, arrestin (9 µM) and varying amount of 08:0 PI(4,5)P_2_ (5 µM) were incubated in buffer containing 20 mM HEPES pH 7.4, 100 mM NaCl, 0.004% LMNG, 0.0004% CHS. Using a Prominence-i LC autosampler (Shimadzu), 10 µL was injected onto an ENrich size-exclusion chromatography 650 10 × 300 column (Bio-rad) pre-equilibrated in 20 mM HEPES pH 7.4 100 mM NaCl, 0.004 % LMNG, 0.0004% CHS, and run at a flow rate of 0.8 ml/min. Tryptophan fluorescence was monitored at λ (EX) of 280 nm and λ (EM) of 340 nm.

### Fluorescence anisotropy measurements

BODIPY-TMR phosphatidylinositol 4,5-bisphosphate (BODIPY-PIP_2_) (Echelon Biosciences) was dissolved to a stock concentration of 1 mM in 50 mM HEPES pH 7.4 and used at a final concentration of 4 nM in the assay. For binding measurements, a two-fold dilution series was made from a stock of βarr1 variant to yield fourteen samples with final concentrations ranging from 150 µM to 0.02 µM. A control sample containing buffer only was included to measure the free anisotropy of BODIPY-PIP_2_. After mixing the BODIPY-PIP_2_ with arrestin or buffer, samples were incubated for 1 h at room temperature prior to measurements. Samples were measured in five 20 µL replicates in a black 384-well plate on a Tecan Infinite M1000 (Tecan Life Sciences), using an excitation wavelength of 530 nm, an emission wavelength of 573 nm and bandwidths of 5 nm. The data was fit using a one-site total binding model as described previously^25^.

### Bulk FRET measurements

Bulk FRET measurements were performed on either a Fluorolog instrument (Horiba) using FluorEssence v3.8 software and operating in photon-counting mode, or a Tecan Infinite M1000 PRO multimodal microplate reader (Tecan), as previously described^25^. The FRET efficiencies of AF488-AT647N- and AF488-AT643-labeled βarr1s were calculated as *I*_A_/(*I*_D_ + *I*_A_), where *I*_D_ and *I*_A_ are the maximum intensities of the donor and acceptor emission peaks, respectively.

### HDX-MS experiments

For HDX-MS experiments, a working sample of protein and diC_8_-PI(4,5)P_2_ was prepared by first mixing βarr1 and PIP_2_ at a 1:10 molar ratio with concentration of βarr1 >100 μM in 20 mM HEPES pH 7.4 100 mM NaCl, 0.004 % LMNG, 0.0004% CHS. This mixture was then diluted to 20.0 μM βarr1 and 200.0 μM PIP_2_ using the same buffer and used directly.

For HDX-MS experiments, the labeling buffer and quench buffer recipes are as follows: 20 mM HEPES, 100 mM NaCl, pD equal to 7.4 in D_2_O, and 2 M guanidine hydrochloride, 100 mM citric acid, 2.3 pH in H_2_O. The pH of the labeling buffer was measured with a pH meter and corrected by pD (pD = pH + 0.4). 4 µl of sample (protein alone or protein + PIP_2_) was diluted 1/7 with labeling buffer and incubated in D_2_O buffer at 10 °C for varying amounts of time in triplicate. Non-deuterated controls were prepared in a similar manner except H_2_O buffer was used in the labeling step. The labeling was quenched by adding an equal volume of quench buffer for 180 seconds and the quenched samples were immediately subjected to online digestion. LC/MS bottom-up HDX was performed using a Thermo Scientific™ Ultimate™ 3000 UHPLC system and Thermo Scientific^TM^ Orbitrap Eclipse^TM^ Tribrid^TM^ mass spectrometer. Samples were digested with a Pepsin/FPXIII 1:1 dual-protease column (NovaBioAssays) at 8°C and desalted using a 1.0 mm x 5.0 mm, 5.0 µm trap cartridge for 3 minutes in total. Peptides were separated on a Thermo Scientific™ Hypersil Gold^TM^, 50x1 mm, 1.9 µm, C18 column with an elution gradient of 10% to 35%B (Buffer A: water + 0.1% formic acid; Buffer B: acetonitrile + 0.1% formic acid) for 10 minutes at 40 µL/min flow rate. To limit carry-over, a pepsin wash was added between runs. The quenching, trapping, and separation steps were performed at near 0 °C to limit back-exchange. Labeling, quenching, and online digestion were performed in a fully automated CHRONECT HDX workstation (Trajan). Both protein and protein-ligand complex samples were analyzed in triplicate for statistical evaluation.

Prior to HDX-MS experiments, a library was created for WT βarr1 with separate MS/MS measurements of non-deuterated samples. Undeuterated WT digested peptides were identified on the orbitrap mass spectrometer using the same LC gradient as the HDX-MS experiment with a combination of data-dependent and targeted HCD-MS2 acquisition. Altogether, 223 peptide assignments were confirmed for WT βarr1, giving 96% sequence coverage. Similarly, we obtained >95% sequence coverage for both R393Q and 3A. HDX experiments had the following βarr1 coverage 90% (WT ± PIP_2_), 88% (WT vs 3A), 95% (WT vs R393Q), 96% (R393Q ± PIP_2_), and 92% (3A ± PIP_2_). MS data files were processed using the HDExaminer software (Trajan) with the WT peptide database. Following the automated HDX-MS analysis, manual verification of the data and correction were performed. Upon completion of the data review, a single charge state with high-quality spectra for all replicates across all HDX labeling times was chosen to represent each peptide. Differential HDX data were tested for statistical significance using the hybrid significance testing criteria method^84^ with an in-house Matlab script. The significant differences observed at each residue were used to map HDX consensus effects (based on overlapping peptides) onto the model of PBD ID: 1G4M using ChimeraX. Residue level data analysis was performed using built-in functions in BioPharma Finder 5.1 software^54^. The used parameters are as follows: 200 simulations, 200 solutions, Chi^2^ increase by the larger of smooth absolute = 0, smooth relative % = 2, differential absolute = 0, differential relative 2%.

### Surface plasmon resonance measurements

SPR experiments were performed using a GE Biacore T100 instrument. Approximately 300-400 resonance units (RU) of FPLC-purified biotinylated βarrs in HBS-P+ Buffer (Cytiva) were captured on an SA-chip (Cytiva), including a reference channel for online background subtraction of bulk solution refractive index and for evaluation of non-specific binding of analyte to the chip surface (Biacore T100 Control Software; Cytiva). All measurements were performed with 2-fold serial dilutions using 60 s or 120 s association followed by a dissociation time of more than 240 s at 25°C with a flow rate of at least 30 ml min^-1^. Measurements of titrations at equilibrium were used to determine K_d_ values using Biacore Analysis Software (v.2.0.4, Cytiva) and fits to a total binding model were performed in GraphPad Prism 9. Regeneration was performed by 2 injections of 2 M MgCl_2_ for 10 s at 50 ml min^-1^ flow rate, resulting in a complete return to baseline in all cases.

### TIRF-based single-molecule FRET imaging

Microfluidic imaging chambers were prepared by overlaying passivating quartz glass slides with passivated coverslips separated by double-sided tape. Cleaned slides/coverslips were passivated as follows: first glass was treated with 3-aminopropyltriethoxysilane in acetone, followed by treatment with a mixture of 5,000 MW polyethylene glycol (PEG-5k) and biotin-PEG-5k succinimidyl valerates (SVA). For direct immobilization of βarr1 variants, the imaging surface was treated with 1 µM NeutrAvidin, followed by ∼250 pM biotinylated βarr1. After 5 minutes, unbound βarr1 protein was flushed out with an imaging buffer (20 mM HEPES-KOH pH 7.4, 100 mM NaCl, 2 mM Trolox).

smFRET imaging experiments were performed at 21°C with a custom-built prism TIRF microscope. Immobilized proteins were illuminated with a 488 nm solid-state laser (Obis, Coherent). Fluorescence emission from AlexaFluor488 and ATTO647N was collected by a 60X, Plan apo 1.27 NA WI objective (Nikon), passed through a 496LP filter (Semrock) to remove excitation light, reflected with a ZT488rdc UF1 dichroic (Chroma), spectrally split in a TwinCam Device (Cairn) with a ZT633rdc-UF2 dichroic filter (Chroma), and projected onto two synchronized iXon Ultra 888 EMCCD cameras (Andor) with 1x1 pixel binning and operating at - 60 °C with between 750 and 1000 EM gain. The donor emission channel was further cleaned with an ET535/70 bandpass filter (Chroma). Recordings were made either at 100 ms time resolution (10 Hz) or 10 ms time resolution (100 Hz) and are indicated for each experiment. Instrument control was performed with Andor Solis (Andor) and Obis connection (Coherent).

### Single-molecule data analysis

The donor and acceptor fluorescence time trajectories from each molecule were extracted and analyzed using custom software implemented in MATLAB, made available as a Github repository: https://github.com/JonathanDeutsch/smFRET.ai. FRET trajectories were calculated as (1 + (*I*_D_/*I*_A_))^-1^, where *I*_D_ and *I*_A_ are the background-corrected donor and acceptor fluorescence intensities at each frame. Since the fluorophores are spectrally well-resolved and of similar brightnesses, additional corrections were not applied to retain the original noise characteristics for kinetic analysis. FRET traces with simultaneous donor-acceptor photobleaching were selected for further analysis using manual filtering. Population histograms of the selected traces were summed over the first 30 or 60 frames for data recorded at 100 ms and 10 ms camera integration times, respectively. Gaussian mixture models (GMMs) were fit to selected traces directly to obtain the average FRET and occupancy of each state in the model. State centers obtained from fitting basal βarr1 imaged at 10 ms were used as initial guesses for the low-, mid- and high-FRET states for all conditions.

To quantify the kinetics of the C-tail, FRET traces were idealized using a hidden Markov model (HMM) as implemented in the vbFRET software^48^, followed by k-means clustering to assign states and to reduce overfitting of fluctuations by the HMM. Transition density plots (TDPs) were extracted from the resulting state sequences using transitions corresponding to ΔFRET_ideal_ ≥ 0.15. The number of states *k* used for each condition was determined based on the lowest value for which a *k* + 1 state model did not meaningfully change the clusters in the TDP. To calculate the mean dwell times for different protein states, survival curves derived from a single concatenated state sequence were fit to stretched or double exponential decay functions using non-linear regression as noted in the figure legends. When a double exponential was used, the mean relaxation is reported as the intensity weighted average. Rates are only reported for two-state systems where they are estimated by the inverse of the mean dwell time of the originating state. Kinetics are reported as the bootstrapped mean ± s.d. of 100 resamples of the viterbi, with resampling probability weighted by the trajectory lengths. Each bootstrap sample has a different random order in which it is concatenated before state assignment, such that variation due to stitching is reflected in the reported errors. Analysis of the selected and idealized traces were performed in Python 3.9.

### Quantification and statistical analysis

Quantification and statistical tools used for each set of experiments in this study are outlined in the relevant method sub-section or figure legends. The t-tests described above assume data are normally distributed. While individual datapoints appear consistent with this assumption, given the small sample size we did not perform explicit tests of normality. For single-molecule experiments, uncertainties were estimated using 100 bootstrap samples of the FRET traces as previously described^85,86^. Error bars and bands denote the 95% confidence interval or standard error of the given parameter obtained from the sampling distribution as specified.

---

**Figure S1.**
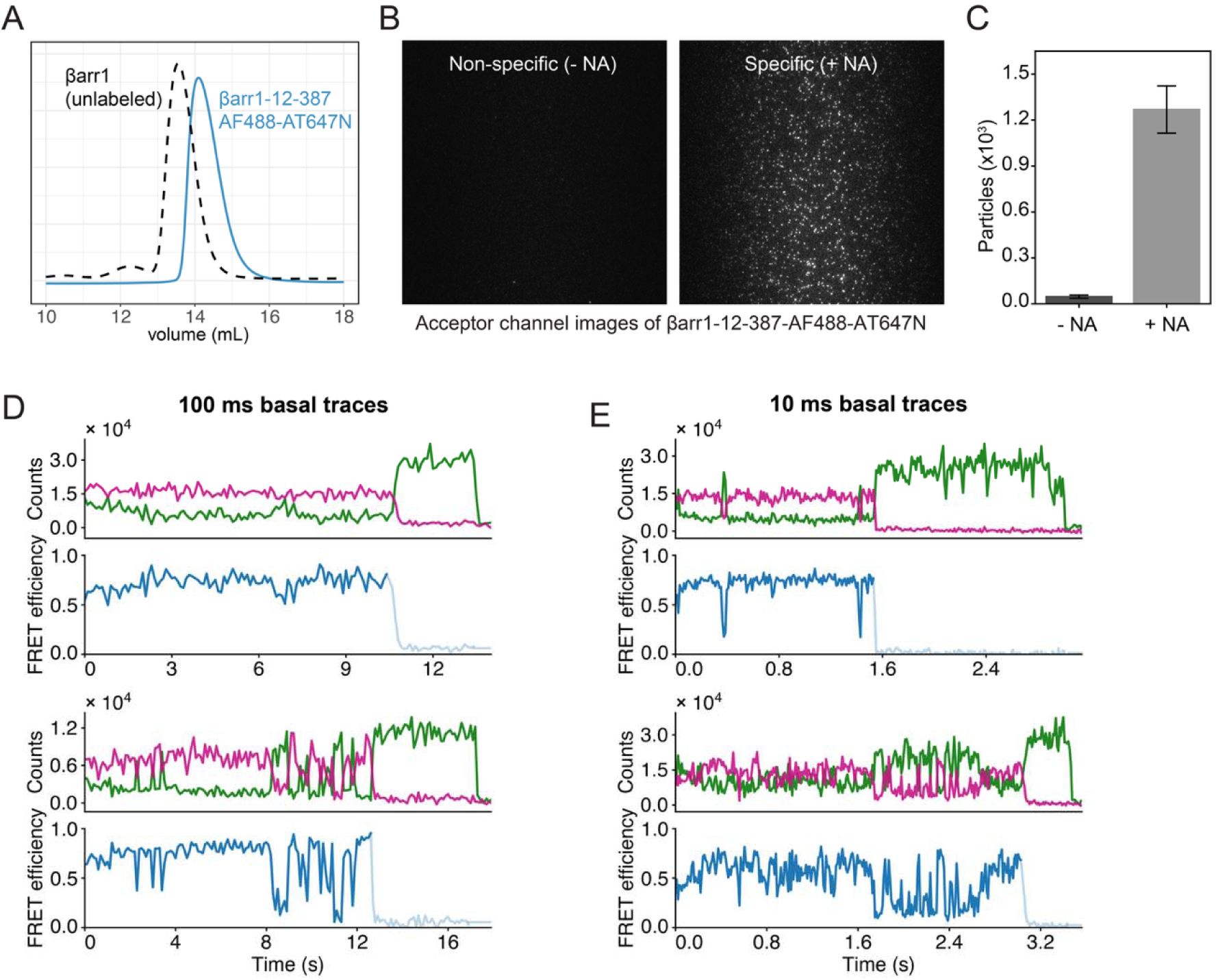
Controls for smFRET imaging of βarr1, related to Figure 2. (A) Size exclusion chromatogram from the purification of unlabeled (black dashed) and fluorophore-conjugated (blue solid) βarr1-12-387. (B and C) (B) Static images of surface-tethered βarr1-biotin conjugate after incubation in the absence (left) and presence (right) of ∼3 μM neutravidin (NA), and (C) histogram of immobilized FRET particles. (D and E) Example single-molecule fluorescence (donor in green; acceptor in magenta) and FRET time traces of surface-tethered βarr1 in the basal state recorded at (D) 100 ms and (E) 10 ms imaging speeds.

**Figure S2.**
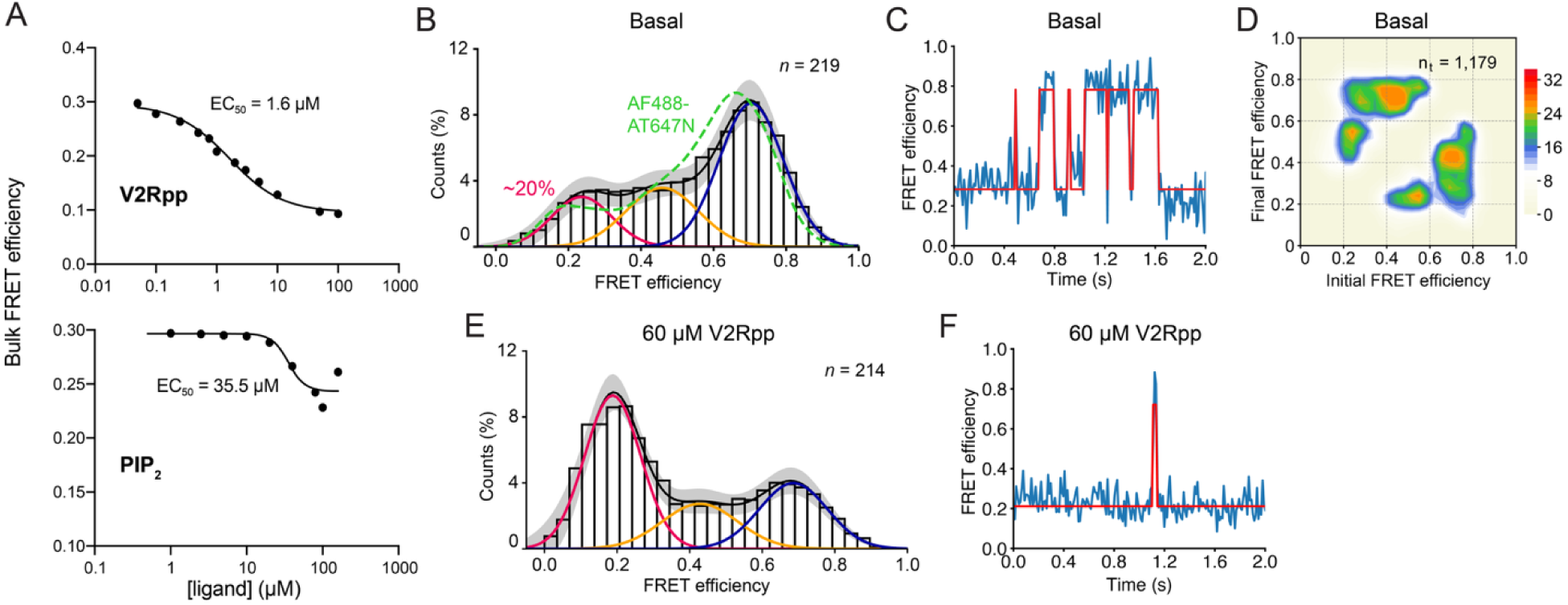
Alternative dye pair for measuring βarr1 C-tail dynamics, related to Figures 2 and 3. (A) Functionality of βarr1-12-387-AF488-AT643 demonstrated from changes in bulk FRET efficiency (symbols) induced by various concentrations of V2Rpp (top) and PIP_2_ (bottom). Lines are fits to dose response curves with EC_50_ values of 1.6 μM and 35.5 μM for V2Rpp and PIP_2_, respectively. (B-D) (B) Population FRET efficiency histogram, (C) example smFRET trace, and (D) transition density plot (scale bar, 10^-3^ transitions per bin per second for *n_t_*total transitions; normalized to Figure 2D) for AF488-AT643-labeled βarr1 imaged in the basal state at 10 ms. (E and F) (E) Population FRET histogram and (F) example smFRET trace in the presence of 60 μM (saturating) V2Rpp. Error bands, 95% c.i. of 100 bootstrap samples of the *n* FRET traces.

**Figure S3.**
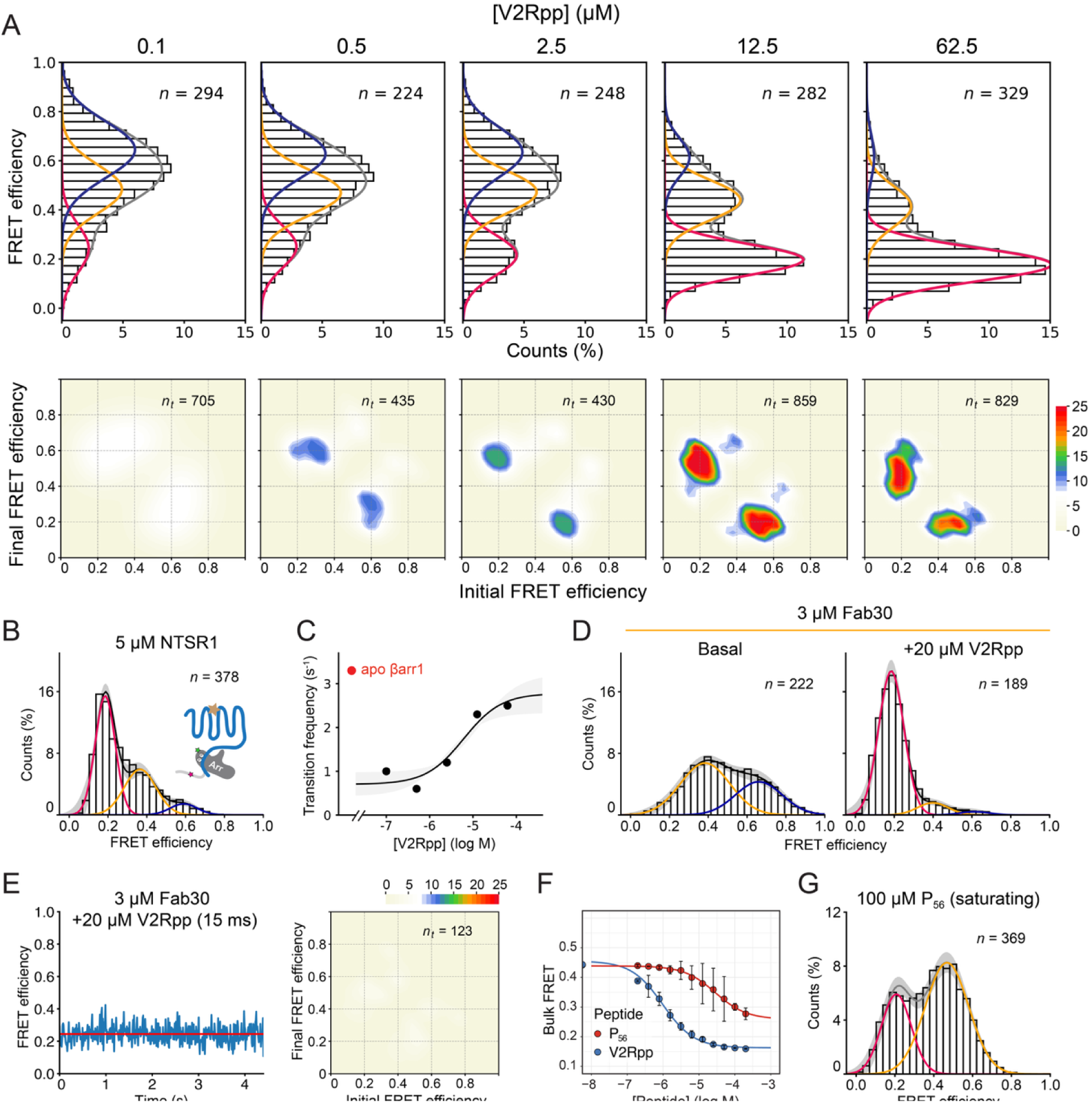
Supporting data for Rp tail mimetics, related to Figure 3. (A) Population smFRET histograms (top; n, number of molecules) and corresponding transition density plots (scale bar, 10^-3^ transitions per bin per second; *n_t_* transitions per condition). (B) Population FRET histogram from imaging the C-tail of surface-tethered βarr1 at 100 ms in the presence of 5 μM NTSR1 bound to neurotensin peptide fragment (NTS8-13). (C) Ensemble average transition frequencies (circles) for V2Rpp titration fit to a dose-response curve (solid line; error band, bootstrapped 95% c.i. of the regression estimate). (D) Population FRET histograms and GMM fits of βarr1 imaged at 10-15 ms camera integration time in the presence of 3 μM Fab30, as well as in the absence (left) and in the presence (right) of 20 μM V2Rpp. (E) Representation FRET trace (left) and transition density plot (right) from experiments imaging of βarr1 in the presence of Fab30 and V2Rpp. (F) Bulk FRET efficiency induced by various concentrations of V2Rpp (blue) and P_56_ (red) phosphopeptides. Lines are fits to dose-response functions (Hill slope = 1.0) with EC_50_ of 2.3 μM for V2Rpp and 32 μM for P_56_, respectively. (G) Population FRET efficiency histogram and GMM fit for βarr1 imaged in the presence of saturating (100 μM) P_56_. Error bands, 95% c.i. from 100 bootstrap samples of the *n* FRET traces per condition.

**Figure S4.**
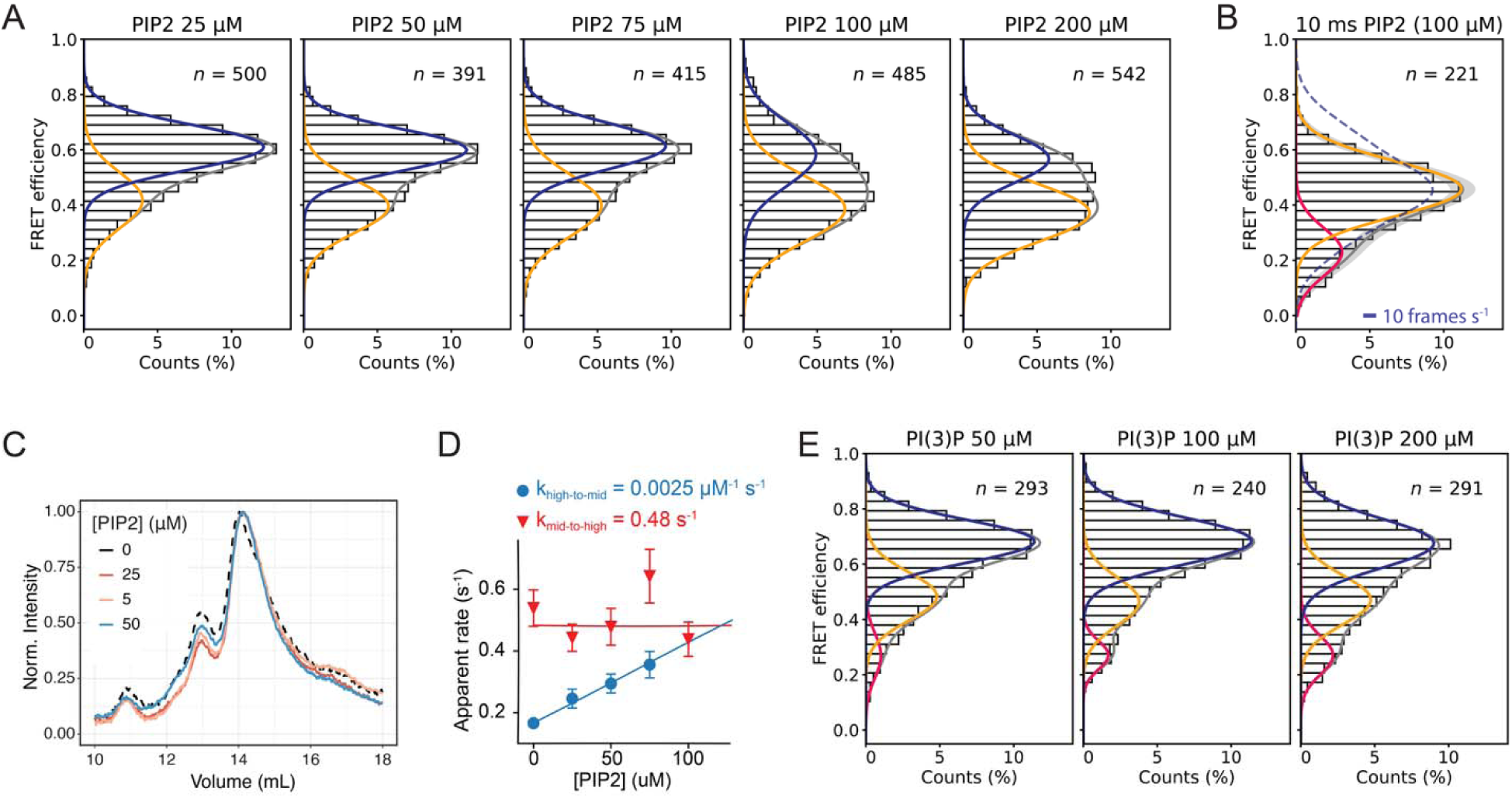
Supporting data for PIP_2_ binding to βarr1, related to Figure 4. (A) Population FRET histograms from experiments imaging βarr1-12-387 in the presence of the indicated concentrations of PIP_2_ at 100 ms time resolution. Cumulative data from each concentration is fit to a two-state GMM (*n* traces per condition). (B) Population FRET histogram and two-state GMM fit for 100 μM PIP_2_ imaged at 10 ms (error bands, bootstrapped 95% c.i.). The dashed line denotes the distribution obtained at 100 ms (10 frames per s). (C) SEC chromatograms of βarr1 in the presence of 0-50 μM PIP_2_ show no effect of the phosphoinositide on oligomerization. (D) Apparent rate constants for PIP_2_ binding (blue) and unbinding (red) of 0.0025 μM^-1^ s^-1^ and 0.48 s^-1^, respectively. Lines are linear fits to the rate constants (symbols) taken as the inverse mean dwell times (Figure 4E). Error bars, mean ± s.d. of 100 bootstrap samples of the idealized FRET traces. (E) Population FRET histograms from experiments with PI(3)P fit to a three-state GMM.

**Figure S5.**
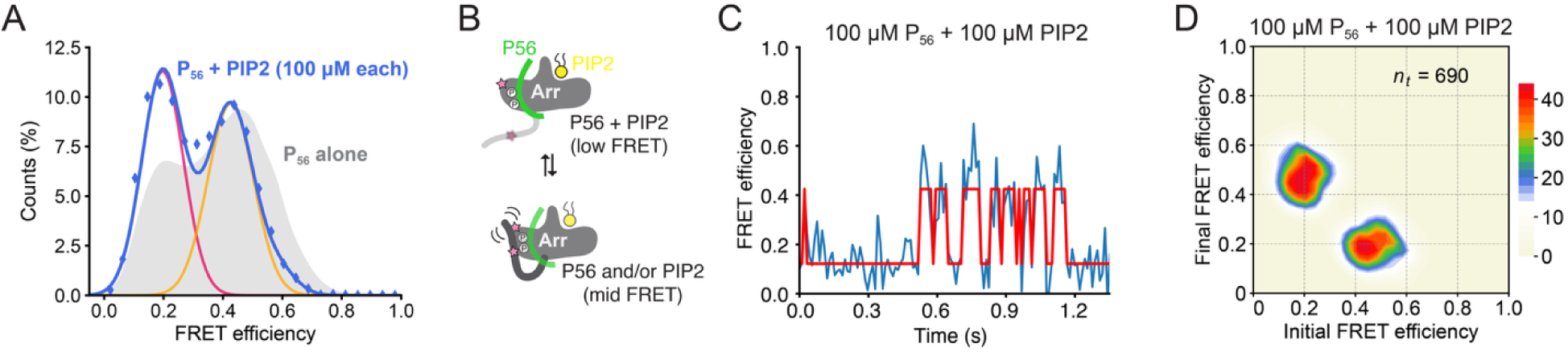
Cooperative interplay between PIP_2_ and P_56_ activation, related to Figures 3 and 4. (A-D) (A) Population FRET histogram (*n* = 278 traces), (B) cartoon representation, (C) example smFRET trace and (D) transition density plot from 10 ms experiments of βarr1-12-387-AF488-AT647N in the presence of both 100 μM PIP_2_ and 100 μM P_56_ phosphopeptide.

**Figure S6.**
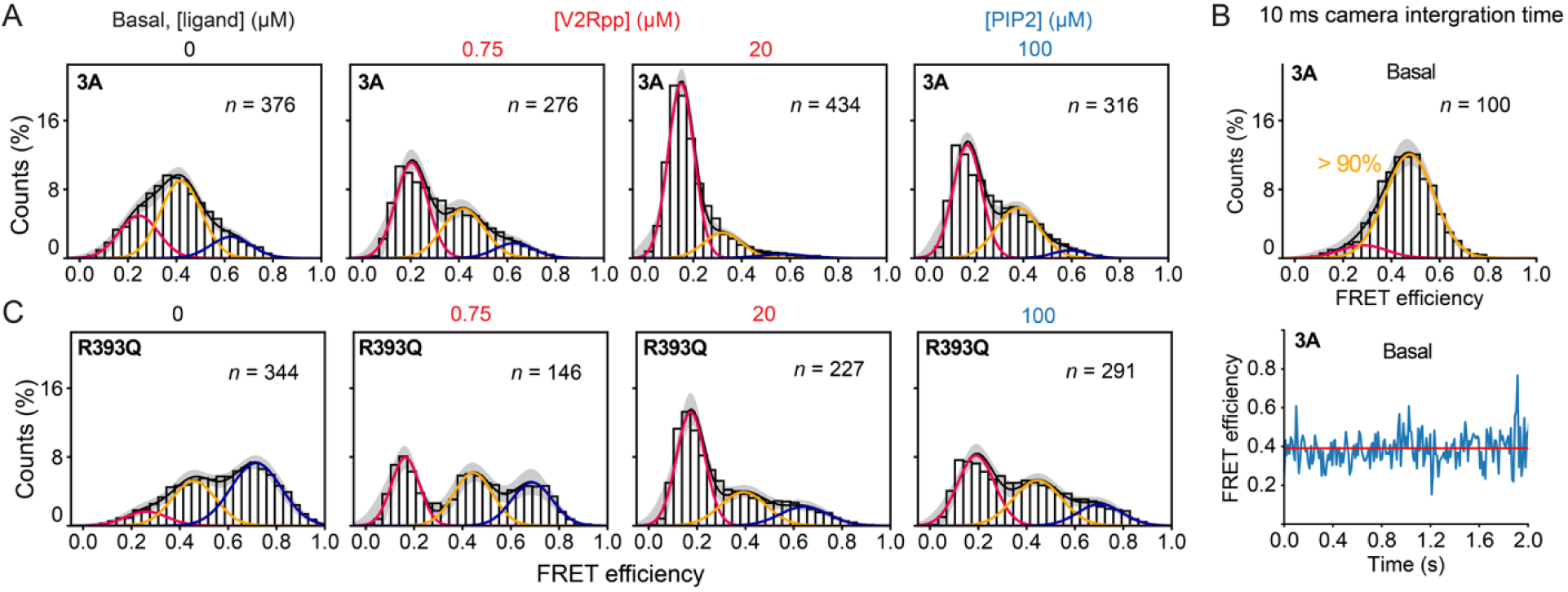
Supporting data for βarr1 C-tail mutants, related to Figure 5. (A) Population FRET histograms with GMM fits from experiments imaging the 3A βarr1 C-tail in the basal state (left), and in the presence of EC_50_ and saturating concentrations of V2Rpp (middle) or 100 μM PIP_2_ (right). Error bands, 95% c.i. of 100 bootstrap samples of the *n* FRET traces per condition, recorded at a 100 ms camera integration time. (B) Imaging 3A βarr1 in the basal state at a 10 ms time resolution showed >90% occupancy of the mid-FRET state (top) and stable mid-FRET dwells (bottom). (C) As in panel A, but with the polar core mutant R393Q βarr1.

**Figure S7.**
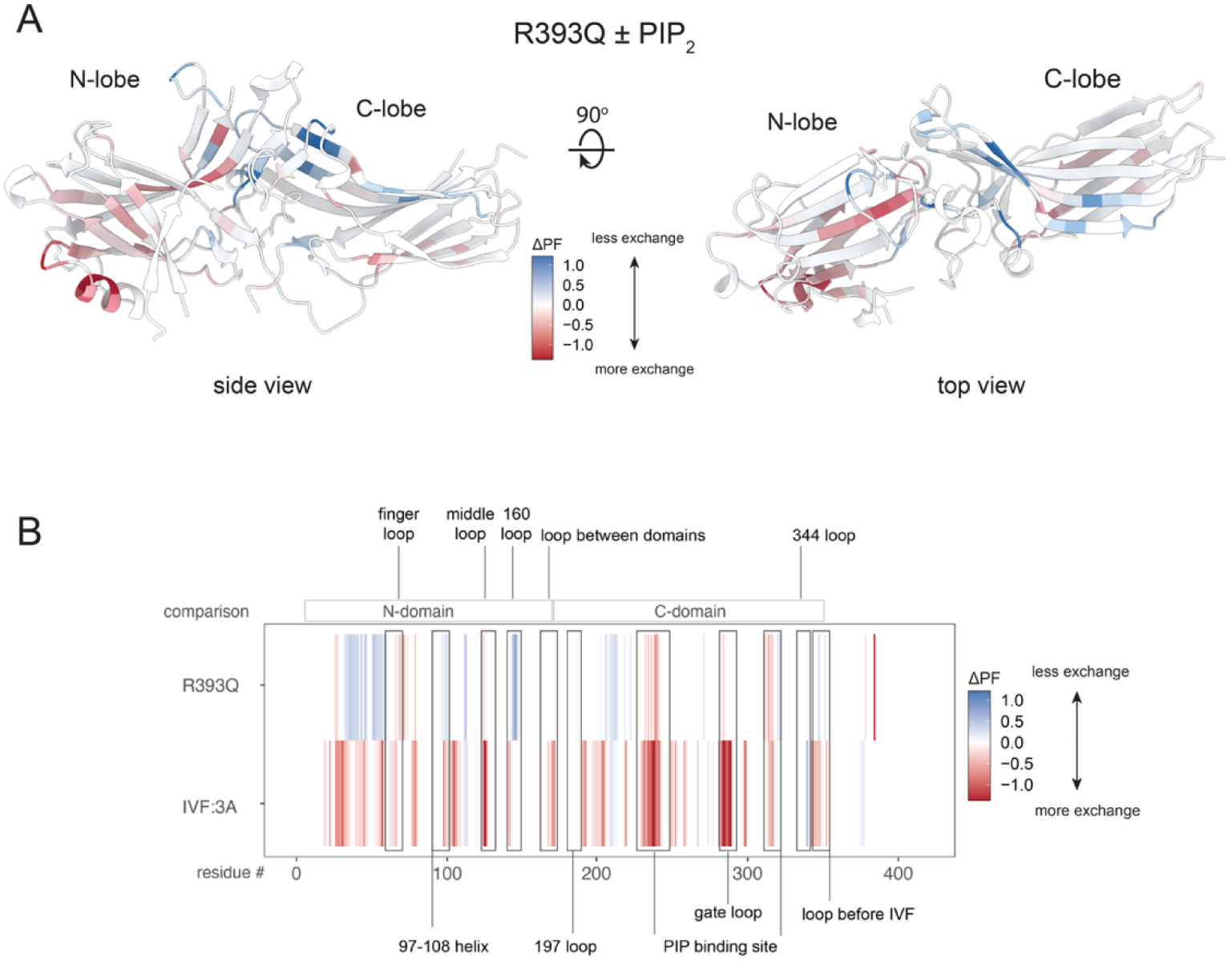
Supporting data for HDX-MS experiments, related to Figure 6. (A) Residues showing significant changes in HDX following the addition of PIP_2_ to R393Q βarr1 (shown on PDB: 1G4M). Scale bar, normalized change in residue-level protection factor: ΔPF = PF_basal_ - PF_PIP2_. (B) Linear map of the change in HDX profile obtained for the R393Q and 3A mutants under basal conditions: ΔPF = PF_MT_ - PF_WT_.

**Figure S8.**
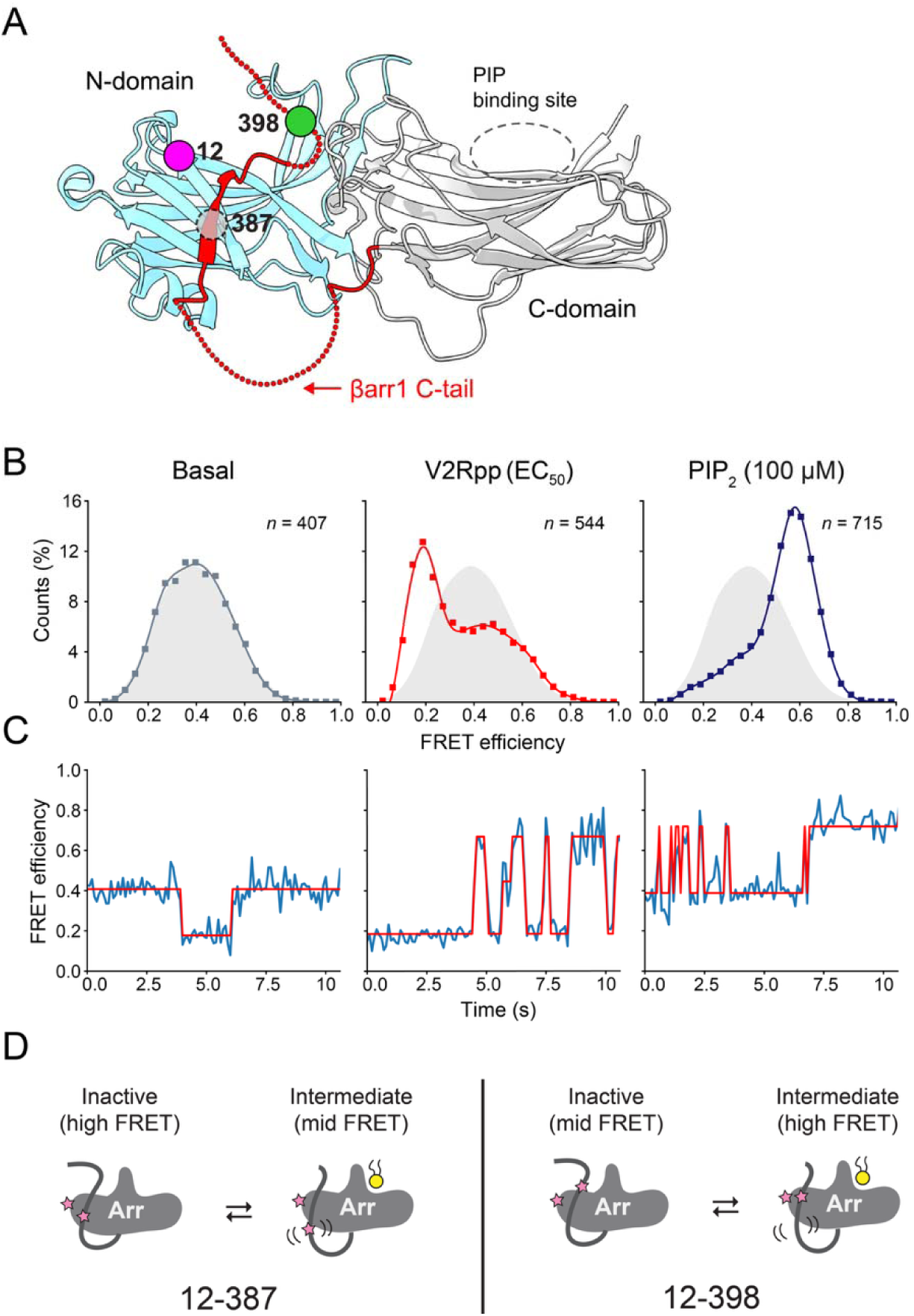
A middle C-tail labeling site shows distinct βarr1 dynamics, related to Figures 4 and 6. (A) Comparison of the middle C-tail labeling site, G398C, to the primary proximal site, V387C (PDB: 1G4M). (B and C) (B) population FRET histograms (symbols) with b-spline fits (lines). Gray density is βarr1 12-398 in the basal state (*n* total molecules) and (C) smFRET trajectories showing representative dynamics from measurements of the βarr1-12-398 middle C-tail sensor in its basal state (left); in the presence of an EC_50_ concentration of V_2_Rpp (0.75 μM; middle); in the presence of saturating 100 μM PIP_2_ (right). (D) Cartoons showing plausible C-tail movements between the two C-tail sensors in response to PIP_2_ binding.

**Figure S9.**
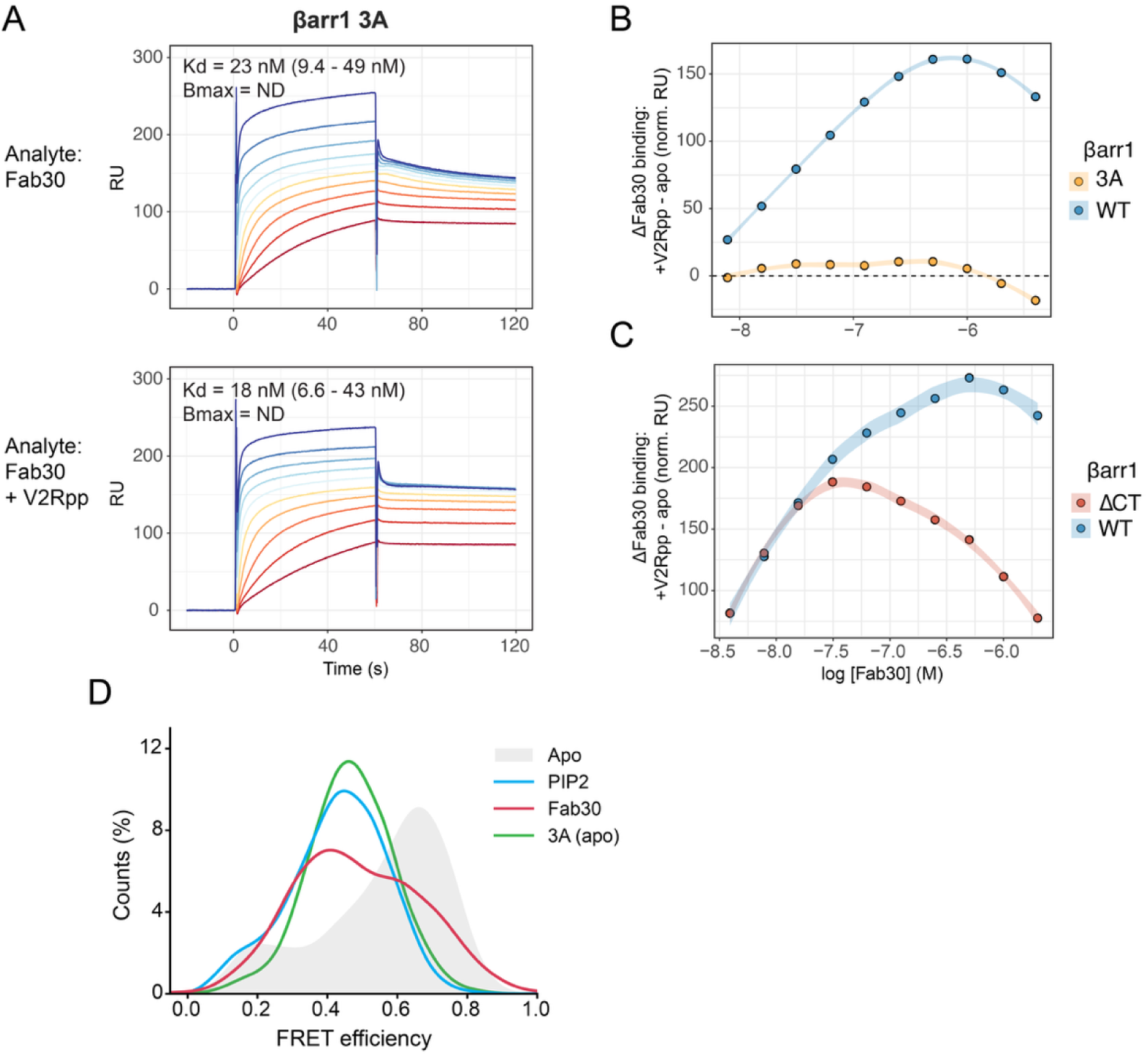
SPR measurements of Fab30 binding to immobilized βarr1, related to Figure 7. (A) SPR sensorgrams of Fab30 binding to 3A βarr1 immobilized via N-terminal biotinylation. Titrations (1:1) of Fab30 ranged from 2 μM to 3.9 nM Fab30. (B and C) Effect of V2Rpp on Fab30 binding by SPR for (B) WT versus 3A and (C) WT versus C-tail truncated (ΔCT) βarr1 constructs. Titrations in the presence (+V2Rpp) or absence (apo) of V2Rpp, as shown in panel A, were used to calculate ΔFab30 binding by subtracting the apo response from that with a fixed concentration of V2Rpp (40 μM). RU values are normalized to correct for the amount of immobilized βarr1; data are from a single independent experiment. Conditions besides 3A are calculated from previously published data for comparison^25^. (E) smFRET histograms of various conditions where βarr1 populates a mid-FRET conformation.

